# Phenotypic plasticity in a novel set of EGFR tyrosine kinase inhibitor-adapted non-small cell lung cancer cell lines

**DOI:** 10.1101/2024.10.29.620819

**Authors:** Tharsagini Nanthaprakash, Campbell W. Gourlay, Ina Oehme, Michelle D. Garrett, Jindrich Cinatl, Mark N. Wass, Martin Michaelis

**Author notes:** Corresponding authors’.

## Abstract

Here, we introduce sublines of the EGFR-mutant non-small cell lung cancer (NSCLC) cell lines HCC827 and HCC4006 adapted to the EGFR kinase inhibitors gefitinib (HCC827^r^GEFI^2µM^, HCC4006^r^GEFI^1µM^), erlotinib (HCC827^r^ERLO^2µM^, HCC4006^r^ERLO^1µM^), and afatinib (HCC827^r^AFA^50nM^, HCC4006^r^AFA^100nM^). All sublines displayed resistance to gefitinib, erlotinib, afatinib, and the third-generation EGFR kinase inhibitor osimertinib that overcomes T790M-mediated resistance. HCC4006^r^ERLO^1µM^ displayed a spindle-like morphology in agreement with previous findings that had detected epithelial-mesenchymal-transition (EMT) in its precursor cell line HCC4006^r^ERLO^0.5µM^. EMT had also been reported for the HCC4006^r^GEFI^1µM^ precursor cell line HCC4006^r^GEFI^0.5µM^ and for HCC4006^r^AFA^100nM^, but the morphologies of HCC4006^r^GEFI^1µM^ or HCC4006^r^AFA^100nM^ did not support this, suggesting plasticity in EMT regulation during the drug adaptation process and in established resistant cell lines. Accordingly, HCC4006^r^ERLO^1µM^ displayed resistance to MEK and AKT inhibitors in contrast to its precursor HCC4006^r^ERLO^0.5µM^. We also detected metabolic plasticity, i.e., a temporary Warburg metabolism, in HCC4006 and HCC827^r^GEFI^2µM^. Response profiles to cytotoxic anti-cancer drugs, kinase inhibitors, and HDAC inhibitors resulted in complex patterns that were specific for each individual subline without obvious overlaps, indicating individual resistance phenotypes. All resistant sublines remained sensitive or displayed collateral sensitivity to at least one of the investigated drugs. In conclusion, the comparison of EGFR kinase-resistant NSCLC sublines with their precursor cell lines that had been previously characterised at a lower resistance level and metabolic investigations indicated phenotypic plasticity during the resistance formation process and in established cell lines. This plasticity may contribute to the well-known variability in cell line phenotypes observed between different laboratories and in intra-laboratory experiments.

## Introduction

Lung cancer is responsible for the highest number of cancer-related deaths, with 85% of cases being non-small cell lung cancer (NSCLC) (1–3). Many NSCLCs are driven by activating epithelial growth factor receptor (EGFR) mutations and are treated by first- (e.g., erlotinib, gefitinib), second- (e.g., afatinib), and/ or third generation (e.g., osimertinib) EGFR tyrosine kinase inhibitors (1–5). However, resistance formation after an initial therapy response is common, and new therapies are needed for NSCLC patients, whose tumours have stopped responding to EGFR tyrosine kinase inhibitor therapy (1–5).

Drug-adapted (cancer) cell lines have been successfully used to identify clinically relevant resistance mechanisms since the 1970s (6–15). Moreover, drug-adapted cancer cell lines enable the detailed analysis of molecular acquired resistance mechanisms and the systematic testing of potential next-line therapies (6,9,14–18).

Here, we introduce a novel set of NSCLC cell lines consisting of HCC827 and HCC4006 and their sublines adapted to gefitinib, erlotinib, and afatinib. The results confirm previous experimental and clinical findings indicating that every resistance formation process follows a unique, unpredictable route. Moreover, they indicate that cancer cell lines are subject to phenotypic plasticity, both during the resistance formation process and as established cell lines.

## Results

### Resistance status, cell morphology, and growth characteristics

All EGFR tyrosine kinase inhibitor-adapted sublines displayed high levels of resistance to the respective inhibitors (Figure 1A). The parental cell lines were sensitive to clinically achievable therapeutic plasma concentrations (C_max_) of the respective EGFR tyrosine kinase inhibitors (19), while the IC_50_ values for all the resistant sublines were above the C_max_ values (Figure 1A, Figure 2A).

**Figure 1.**
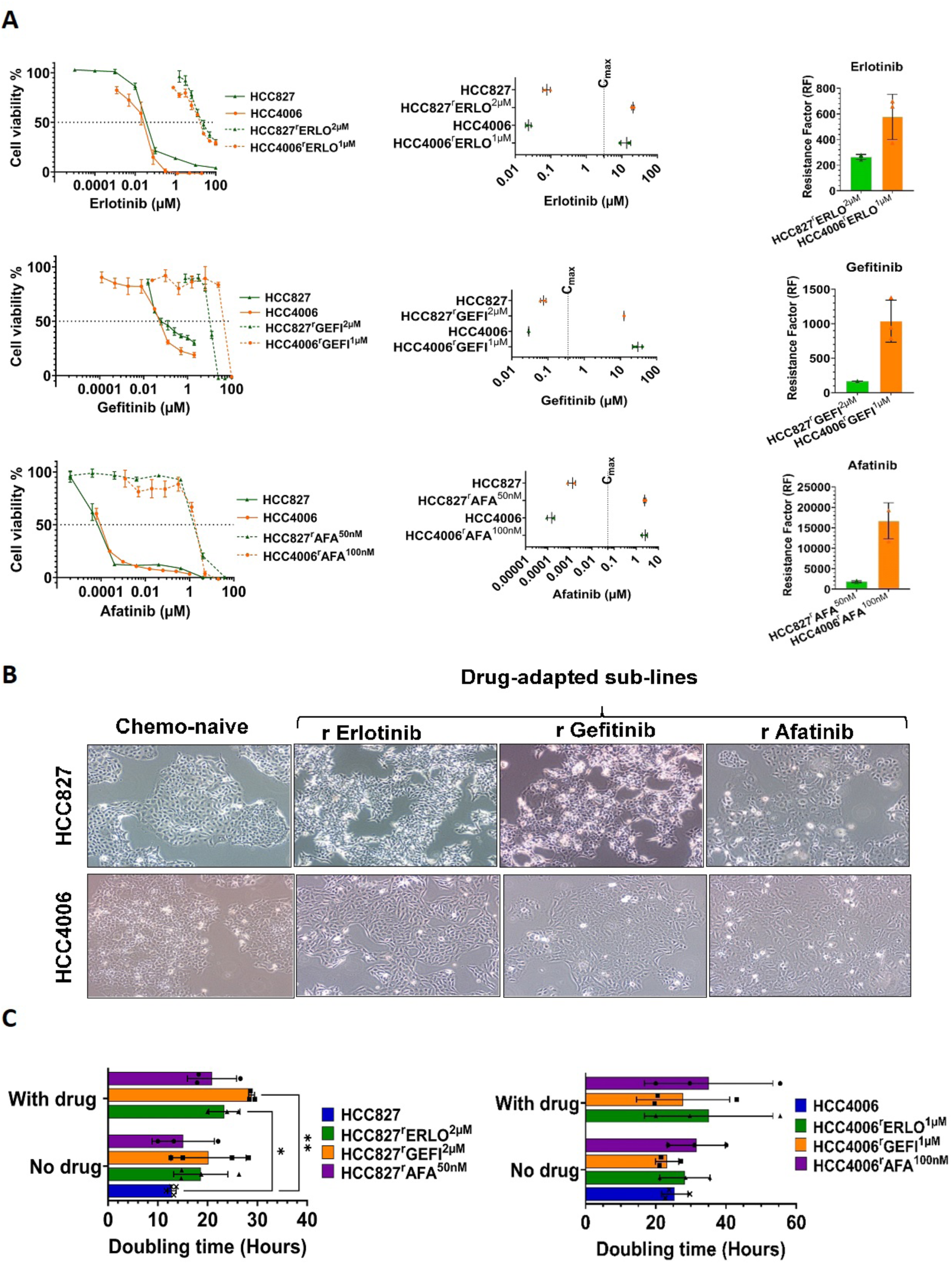
Basic characterization of the EGFR tyrosine kinase inhibitor-adapted HCC827 and HCC4006 sublines. A) Dose response curves, IC_50_ values, and resistance factors (RF, IC_50_ resistant subline/ IC_50_ respective parental cell line) demonstrating that the drug-adapted sublines are resistant to the respective EGFR tyrosine kinase inhibitors relative to the respective parental cell lines. Cell viability was determined by MTT assay after a 120h incubation period. IC_50_ values were calculated using the software Calcusyn (Version 1.1, Biosof 1996). B) Representative images of HCC827, HCC4006, and their EGFR receptor tyrosine kinase-adapted sublines at 40x magnification (Olympus CKX53 inverted microscope, Olympus Life Sciences, UK). C) Doubling times of HCC827, HCC4006, and their EGFR receptor tyrosine kinase-adapted sublines. In the resistant sublines, doubling times were determined in the absence and presence of the respective drugs of adaptation. Differences were analysed for statistical significance (p<0.05) by student’s t-test. *p<0.05, **p<0.01**.

**Figure 2:**
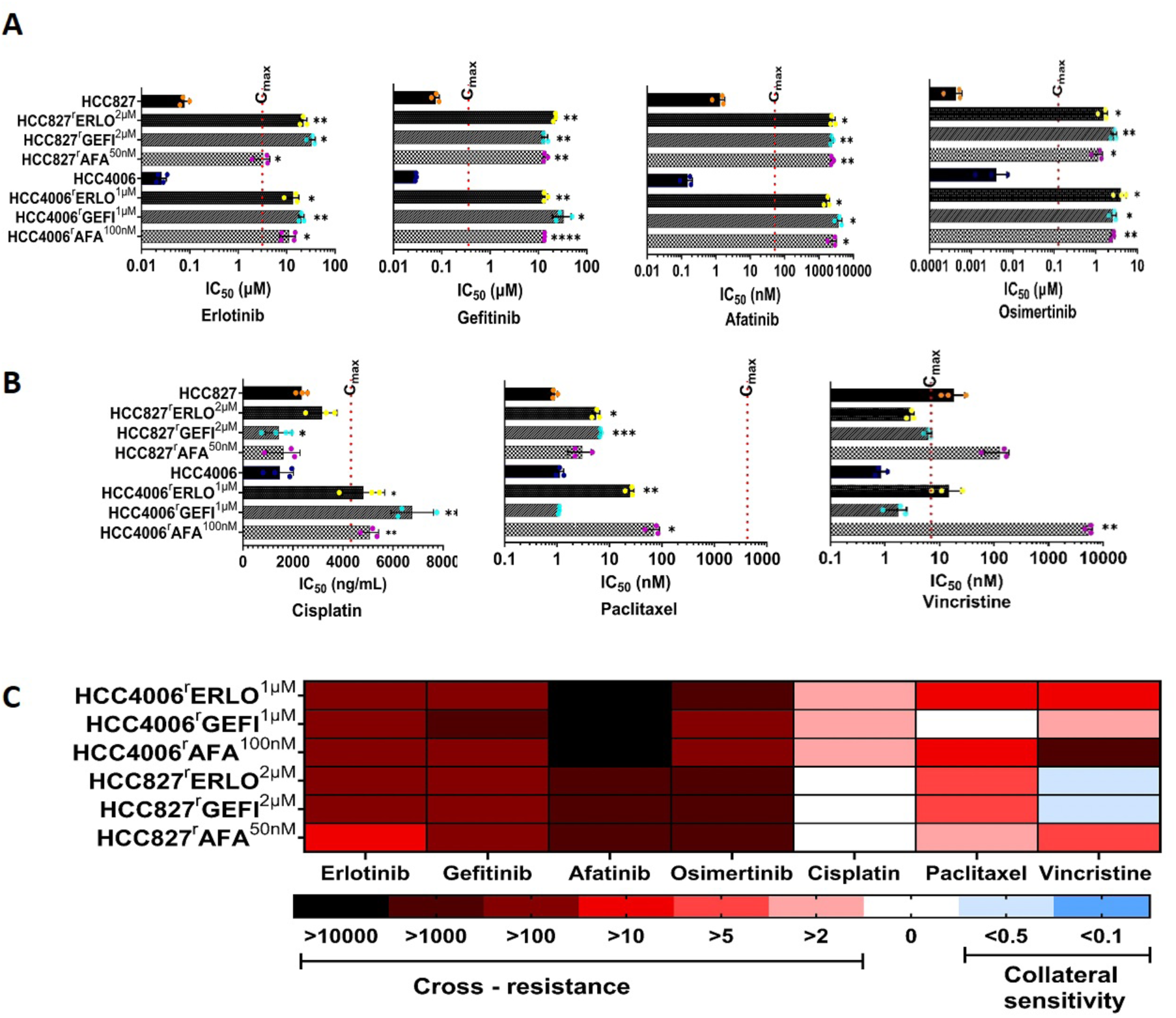
Cell line sensitivity profiles to selected anti-cancer drugs. Cell viability was determined by MTT assay after a 120h incubation period. IC_50_ values were calculated using the software Calcusyn (Version 1.1, Biosof 1996). Values represents means ± S.D. of at least three independent experiments. Dose-response curves are presented in Suppl. Figure 2. Red dotted line represents the clinical maximum plasma concentrations (C_max_) of respective the drugs. Differences were analysed for statistical significance (p<0.05) by student’s t-test. *p<0.05, **p<0.01**, ***p<0.001, ****p<0.0001. A) Sensitivity of HCC827, HCC4006, and their EGFR receptor tyrosine kinase-adapted sublines to different EGFR tyrosine kinase inhibitors. Dose-response curves are presented in Suppl. Figure 1. B) Sensitivity of HCC827, HCC4006, and their EGFR receptor tyrosine kinase-adapted sublines to cisplatin, paclitaxel, and vincristine. Dose-response curves are presented in Suppl. Figure 2. C) Overview heatmap providing the response profiles of the cell lines to the investigated drugs based on the resistance factors (RF, IC_50_ resistant subline/ IC_50_ respective parental cell line). Resistance (red) was defined as an RF>2, comparable sensitivity as RF≤2 and ≥0.5, and increased resistance/ collateral vulnerability as RF<0.5.

Representative images of the cell lines are provided in Figure 1B. HCC827 and HCC827^r^AFA^50nM^ displayed an enhanced adhesion to the cell flask surface compared to HCC827^r^ERLO^2µM^ and HCC827^r^GEFI^2µM^ (and to HCC4006 and its sublines). Due to this, a higher trypsin concentration (0.12% w/v) than our regular one (0.05% w/v) was required for the detachment of HCC827 and HCC827^r^AFA^50nM^ cells. Moreover, HCC827 and HCC827^r^AFA^50nM^ grew in monolayers, while HCC827^r^ERLO^2µM^ and HCC827^r^GEFI^2µM^ formed multilayers.

Among HCC4006 and its sublines, HCC4006^r^ERLO^1µM^ displayed a more spindle-like morphology, while HCC4006^r^GEFI^1µM^ and HCC4006^r^AFA^100nM^ had a wider diameter than HCC4006 (Figure 1B). Moreover, HCC4006 formed multilayers in contrast to its sublines that all grew as monolayers.

The doubling times of the EGFR tyrosine kinase-adapted HCC827 and HCC4006 sublines did not significantly differ from the respective parental cell lines (Figure 1C, Suppl. Table 1). The addition of the respective drugs of adaptation only affected the doubling times of two of the sublines: The doubling time of HCC827^r^GEFI^2µM^ was 28.8 ± 0.5h in the presence of gefitinib 2µM and 20.2 ± 7.3 in the absence of drug. Moreover, the doubling time of HCC827^r^ERLO^2µM^ was 23.4 ± 2.8h in the presence of erlotinib 2µM and 18.6 ± 4.3 in the absence of drug (Figure 1C, Suppl. Table 1).

### EGFR tyrosine kinase inhibitor response profiles

Next, we determined the response profiles of the project cell lines to the clinically approved EGFR tyrosine kinase inhibitors erlotinib, gefitinib, afatinib, and osimertinib. Erlotinib and gefitinib are first-generation EGFR kinase inhibitors. Afatinib is a second-generation EGFR kinase inhibitor. Osimertinib is a third-generation EGFR kinase inhibitor designed to overcome resistance to first- and second-generation EGFR kinase inhibitors that is mediated by T790M EGFR mutations (2,5).

All resistant sublines displayed pronounced resistance against all four EGFR inhibitors (Figure 2A, Suppl. Figure 1, Suppl. Table 2), indicating that EGFR kinase inhibitor resistance is not driven by T790M mutations in the project cell lines. The resistance factors (IC_50_ resistant subline/ IC_50_ respective parental cell line) ranged from 42.2 (HCC827^r^Afa^50nM^, erlotinib) to 24815.1 (HCC4006^r^Gefi^1µM^, afatinib) (Suppl. Table 2).

### Cytotoxic chemotherapy response profiles

Cytotoxic chemotherapy remains an option for the treatment of EGFR-mutant NSCLC after resistance formation to EGFR kinase inhibitors (20–23). Platinum-based therapies are among the most commonly used cytotoxic anti-cancer drugs used at different stages of the treatment of EGFR-mutant NSCLC (5,20,22–25). Moreover, taxanes and vinca alkaloids belong to the chemotherapeutic drug classes that are still investigated for the treatment of EGFR-mutant NSCLC (5,21,25,26).

Here, we investigated the response of the cell lines from our panel to the platinum drug cisplatin, the taxane paclitaxel (stabilising tubulin-binding agent), and the vinca alkaloid vincristine (destabilising tubulin-binding agent) (Figure 2B, Suppl. Figure 2, Suppl. Table 3).

When analysing the resistance profiles, we considered resistance factors (IC_50_ resistant subline/ IC_50_ respective parental cell line) ≥ 2 as cross-resistance, resistance factors < 2 and > 0.5 as similar sensitivity, and resistance factors ≤ 0.5 as increased sensitivity/ collateral vulnerability. The drug response profiles revealed a substantial level of heterogeneity among the EGFR kinase inhibitor-resistant NSCLC sublines (Figure 2C).

All HCC827 sublines displayed similar cisplatin sensitivity as HCC827 with resistance factors ranging from 0.61 (HCC827^r^GEFI^2µM^) to 1.35 (HCC827^r^ERLO^2µM^) (Figure 2B, Suppl. Table 3A). In contrast, all HCC4006 sublines were cross-resistant to cisplatin with resistance factors ranging from 3.26 (HCC4006^r^ERLO^1µM^) to 4.58 (HCC4006^r^GEFI^1µM^) (Figure 2B, Suppl. Table 3B).

All EGFR kinase-resistant sublines, except for HCC4006^r^GEFI^1µM^ (resistance factor 0.96), displayed cross-resistance to paclitaxel (Figure 2B, Suppl. Table 3B). However, the resistance factors varied considerably among the cross-resistant sublines ranging from 3.38 (HCC827^r^AFA^50nM^) to 61.5 (HCC4006^r^AFA^100nM^).

The vincristine response profiles were associated with the highest level of variability (Figure 2B, Suppl. Table 3). All HCC4006 sublines and HCC827^r^AFA^50nM^ displayed cross-resistance with resistance factors ranging from 2.06 (HCC4006^r^GEFI^1µM^) to 5,451 (HCC4006^r^AFA^100nM^), while HCC827^r^ERLO^2µM^ (resistance factor 0.16) and HCC827^r^GEFI^2µM^ (resistance factor 0.33) displayed collateral vulnerability.

### Kinase inhibitor response profiles

In EGFR-mutant NSCLC, oncogenic signalling by constitutive active EGFR is mediated via the RAS/ RAF/ MEK/ ERK (MAPK) and/ or PI3K/ AKT signalling pathways. In agreement, common EGFR kinase inhibitor resistance mechanisms include the reinstatement of MAPK and PI3K/ AKT signalling by mechanisms including MET amplification (2,5,23,27). Hence, we next investigated the effects of MET (cabozantinib), MEK (trametinib), and PI3K (alpelisib, LY294002) inhibitors on the project cell lines. We also included AT13148, an inhibitor of AGC kinases including AKT, ROCK1/2, and p70S6K (28), into our analysis.

Overall, the EGFR tyrosine kinase-adapted NSCLC sublines displayed complex response patterns to the investigated kinase inhibitors (Figure 3, Suppl. Figure 3, Suppl. Table 4). The HCC827 sublines showed similar or increased sensitivity to all kinase inhibitors relative to HCC827. Among the HCC827 sublines, the kinase inhibitor response profiles were more similar between HCC827^r^ERLO^2µM^ (collateral vulnerability to all tested kinase inhibitors except AT13148) and HCC827^r^GEFI^2µM^ (collateral vulnerability to all tested kinase inhibitors) compared to HCC827^r^AFA^50nM^ (collateral vulnerability only to cabozantinib) (Figure 3, Suppl. Table 4).

**Figure 3.**
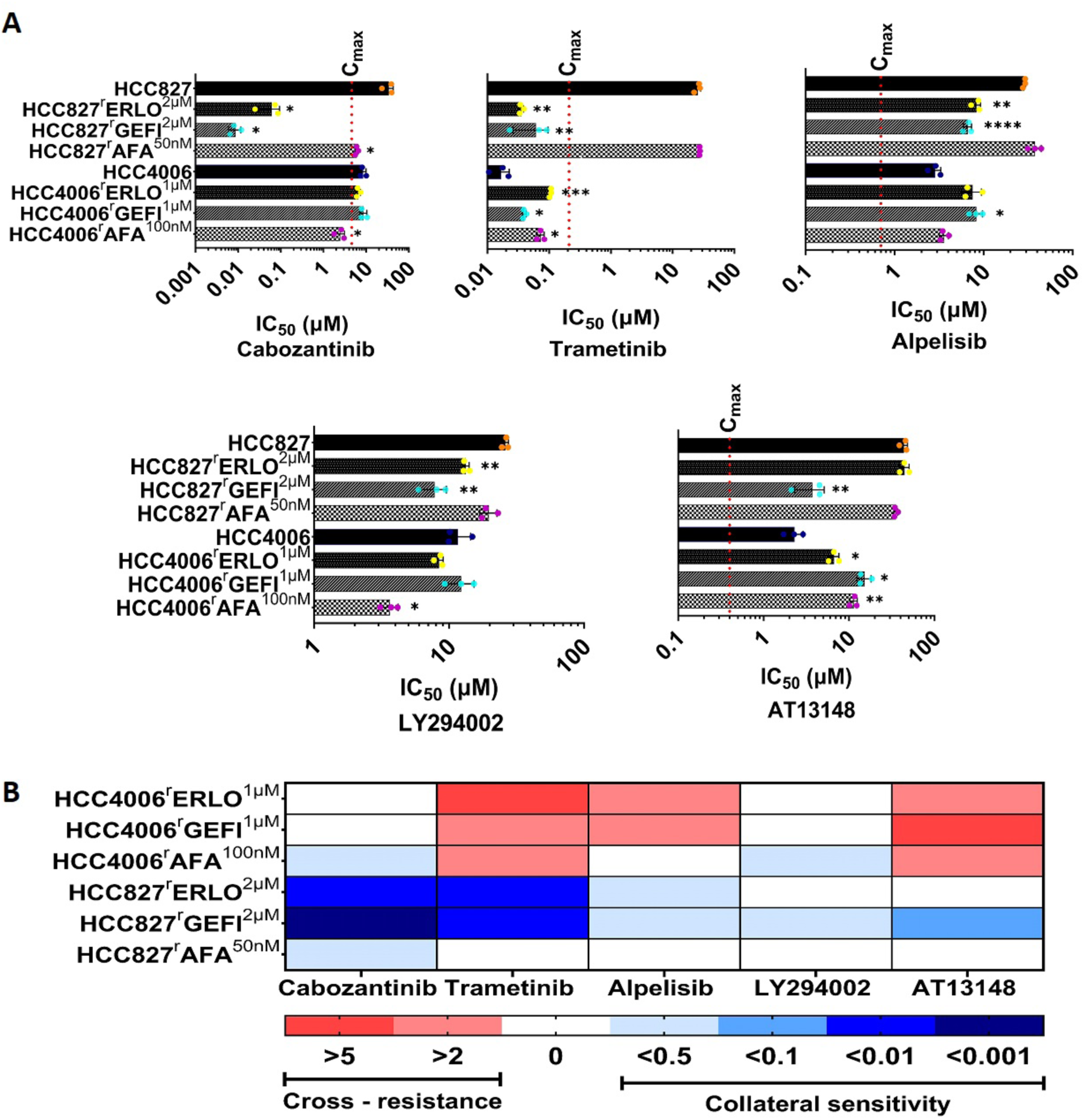
Cell line sensitivity profiles to selected kinase inhibitors. Cell viability was determined by MTT assay after a 120h incubation period. IC_50_ values were calculated using the software Calcusyn (Version 1.1, Biosof 1996). Values represents means ± S.D. of at least three independent experiments. Dose-response curves are presented in Suppl. Figure 3. Red dotted line represents the clinical maximum plasma concentrations (C_max_) of respective the drugs. Differences were analysed for statistical significance (p<0.05) by student’s t-test. *p<0.05, **p<0.01**, ***p<0.001, ****p<0.0001. A) Sensitivity of HCC827, HCC4006, and their EGFR receptor tyrosine kinase-adapted sublines to different kinase inhibitors. Dose-response curves are presented in Suppl. Figure 3. B) Overview heatmap providing the response profiles of the cell lines to the investigated drugs based on the resistance factors (RF, IC_50_ resistant subline/ IC_50_ respective parental cell line). Resistance (red) was defined as an RF>2, comparable sensitivity as RF≤2 and ≥0.5, and increased resistance/ collateral vulnerability as RF<0.5.

In contrast to the HCC827 sublines, the HCC4006 sublines were cross-resistant to a number of kinase inhibitors (Figure 3, Suppl. Table 4). Again, the erlotinib- and gefitinib-adapted sublines displayed a higher level of similarity versus the afatinib-resistant subline. HCC4006^r^ERLO^1µM^ and HCC4006^r^GEFI^1µM^ were both cross-resistant to trametinib, alpelisib, and AT13148 and similarly sensitive as HCC4006 to cabozantinib and LY294002. HCC4006^r^AFA^100nM^ showed collateral vulnerability to cabozantinib and LY294002, cross-resistance to trametinib and AT13148, and similar sensitivity as HCC4006 to alpelisib (Figure 3, Suppl. Table 4B). It remains unclear to which extent the differences in the kinase inhibitor response profiles between the HCC827- and HCC4006-sublines are the consequence of the different cellular backgrounds and/ or of chance events during the adaptation process.

It is difficult to draw conclusions from the kinase inhibitor profiles on whether the resistance formation process may have resulted in a dependence on certain signalling pathways in the sublines. This is most obviously demonstrated by the inconsistent responses of the project cell lines to alpelisib, LY294002, and AT13148 that all target PI3K/ AKT signalling (Figure 3, Suppl. Table 4). Only HCC827^r^GEFI^2µM^ displayed a consistent response (collateral vulnerability) to all three compounds, which might indicate an increased dependence on PI3K/ AKT signalling. However, HCC827^r^GEFI^2µM^ also showed increased sensitivity to the MET inhibitor cabozantinib and the MEK inhibitor trametinib (Figure 3, Suppl. Table 4A), which suggests multiple resistance mechanisms.

Next, we treated the project cell lines with the EGFR kinase inhibitors in the presence of IC_25_ and IC_50_ concentrations of the kinase inhibitors (Suppl. Table 5) to see whether kinase inhibitors may re-sensitise the sublines to the respective EGFR kinase inhibitors (Figure 4, Suppl. Figure 4, Suppl. Table 6). In most cell lines the kinase inhibitors had no or only modest (cabozantinib/ HCC4006^r^GEFI^1µM^, trametinib/ HCC827^r^ERLO^2µM^, trametinib/ HCC827^r^AFA^50nM^) effects on the efficacy of the EGFR kinase inhibitors the sublines had been adapted to (Figure 4, Suppl. Table 6).

**Figure 4.**
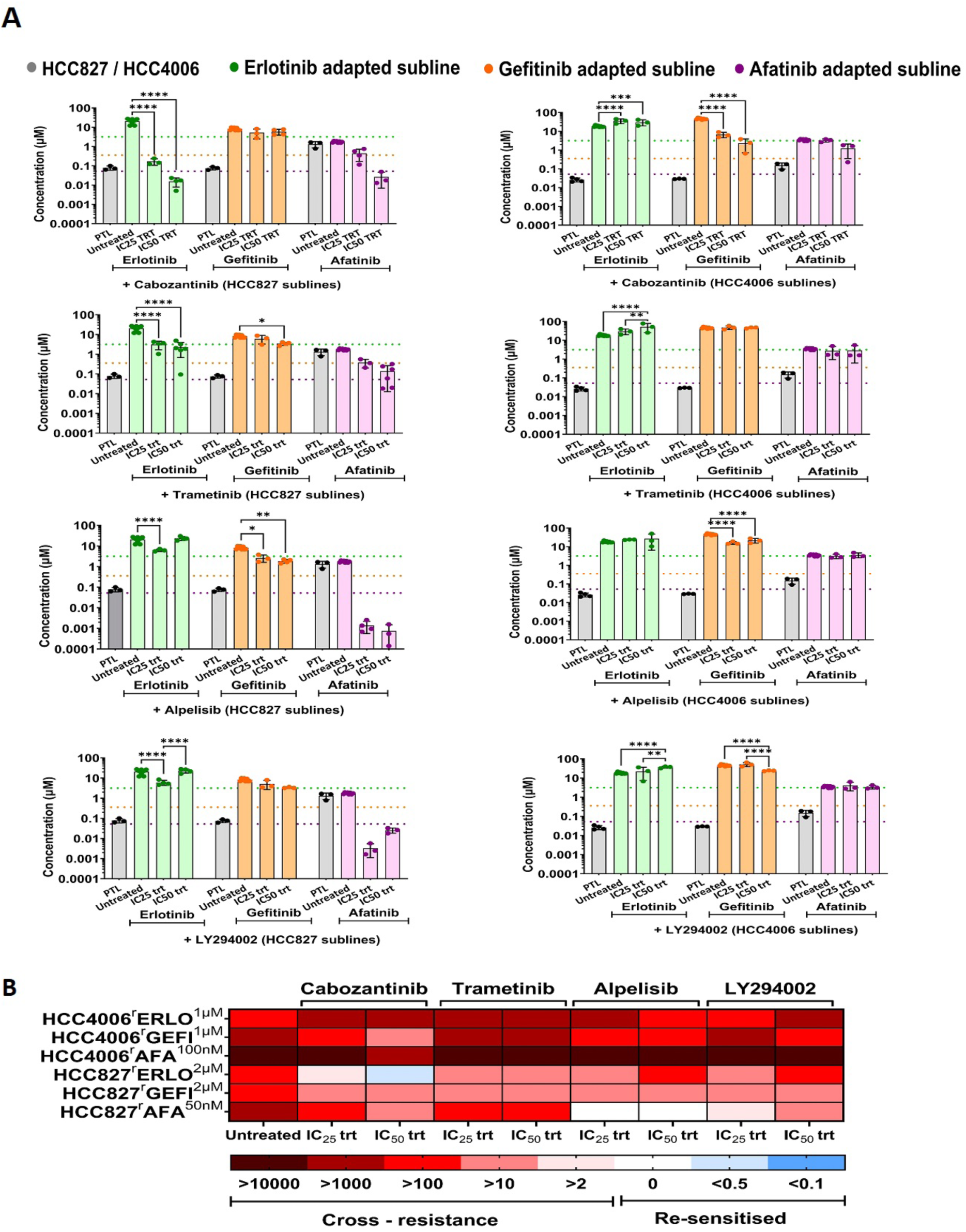
Effect of kinase inhibitors on the sensitivity of EGFR tyrosine kinase inhibitor-adapted cell lines to the respective drugs of adaptation. Cell viability was determined by MTT assay after a 120h incubation period. IC_50_ values were calculated using the software Calcusyn (Version 1.1, Biosof 1996). Values represents means ± S.D. of at least three independent experiments. Dose-response curves are presented in Suppl. Figure 4. Differences were analysed for statistical significance (p<0.05) by Two-way ANOVA with post-hoc Tukey’s multiple comparison test. *p<0.05, **p<0.01**, ***p<0.001, ****p<0.0001. A) Effect of the kinase inhibitors IC_25_ and IC_50_ concentrations on the sensitivity of the EGFR tyrosine kinase inhibitor-resistant sublines to the respective drugs of adaptation. B) Overview heatmap providing the response profiles of the cell lines to the investigated drugs based on the relative resistance (IC_50_ without kinase inhibitor / IC_50_ with kinase inhibitor). Increased resistance (red) was defined as a relative resistance >2, comparable sensitivity as relative resistance ≤2 and ≥0.5, and re-sensitisation as relative resistance <0.5.

Cabozantinib exhibited slightly more pronounced effects on the afatinib activity in HCC827^r^AFA^50nM^ and LY294002 on the afatinib activity in HCC827^r^AFA^50nM^. Re-sensitisation to the level of the parental cell line was only achieved by cabozantinib in HCC827^r^ERLO^2µM^ and by alpelisib in HCC827^r^AFA^50nM^ (Figure 4, Suppl. Table 6). Again, it is difficult to draw meaningful mechanistic conclusions from these results, not least because the PI3K inhibitors alpelisib and LY294002 differed in their activities.

Off-target resistance mechanisms that interfere with compound transport into or out of cancer cells can mediate resistance on their own or in combination with on-target resistance mechanisms that directly affect the drug target and the related signalling pathways (29). In this context, erlotinib, gefitinib, and afatinib are known to be substrates of the ATP-binding cassette transporter ABCB1 (also known as P-glycoprotein or MDR1) (30), a major transporter involved in drug resistance in cancer (29). ABCB1 can also mediate acquired EGFR tyrosine kinase inhibitor resistance (31,32). However, using the ABCB1 substrate vincristine (33) in combination with the specific third-generation ABCB1 inhibitor zosuquidar (33) as previously described (34,35), indicated that only HCC4006^r^ERLO^1µM^ displayed an ABCB1-mediated resistance phenotype (Suppl. Figure 5). This indicates that ABCB1-mediated effects have limited impact on the tyrosine kinase inhibitor response profiles in our panel of EGFR tyrosine kinase-adapted NSCLC sublines.

Notably, other ABC transporters may also be involved in EGFR kinase inhibitor resistance (30,33). Moreover, kinase inhibitors including those that we used in our study are known to display complex interactions with a range of kinases in addition to the desired ones (36–38). Hence, we can conclude that each of the EGFR kinase inhibitor-adapted cell lines has developed a unique phenotype, but future research will have to elucidate the exact resistance mechanisms in each cell line in detail.

### Role of histone deacetylases (HDACs) in EGFR kinase resistance

Changes in histone deacetylase (HDAC) regulation contribute to cancer formation and progression and can also contribute to cancer cell resistance to anti-cancer drugs, including EGFR kinase inhibitors (39–41). The role of HDACs in cancer has resulted in the design of numerous HDAC inhibitors (42). While the first compounds typically were pan-HDAC inhibitors, there has recently been more focus on the development of isotype-specific HDAC inhibitors (13,42–45).

Here, we characterised our cell line panel using a set of 15 HDAC inhibitors, each at a concentration selected to interfere specifically with one or a limited number of HDACs (Suppl. Table 7). HCC827 displayed a generally lower sensitivity to HDAC inhibitors than HCC4006 (Figure 5, Suppl. Table 8). Some resistant sublines displayed an increased sensitivity to certain HDAC inhibitors, in particular HCC827^r^GEFI^2µM^ to apicidin (Figure 5, Suppl. Table 8). However, other HDAC inhibitors that like apicidin inhibit HDAC1, HDAC2, and HDAC3, did not exert comparable effects. Hence, no general mechanistic insights can be drawn from the HDAC inhibitor data. Nevertheless, the findings show that resistance formation to EGFR tyrosine kinase inhibitors can be associated with increased sensitivity to certain HDAC inhibitors. Further research will be needed to investigate the role of HDACs in acquired EGFR kinase inhibitor resistance in more detail. Nevertheless, the findings further confirm that each EGFR kinase-resistant subline has developed an individual phenotype.

**Figure 5.**
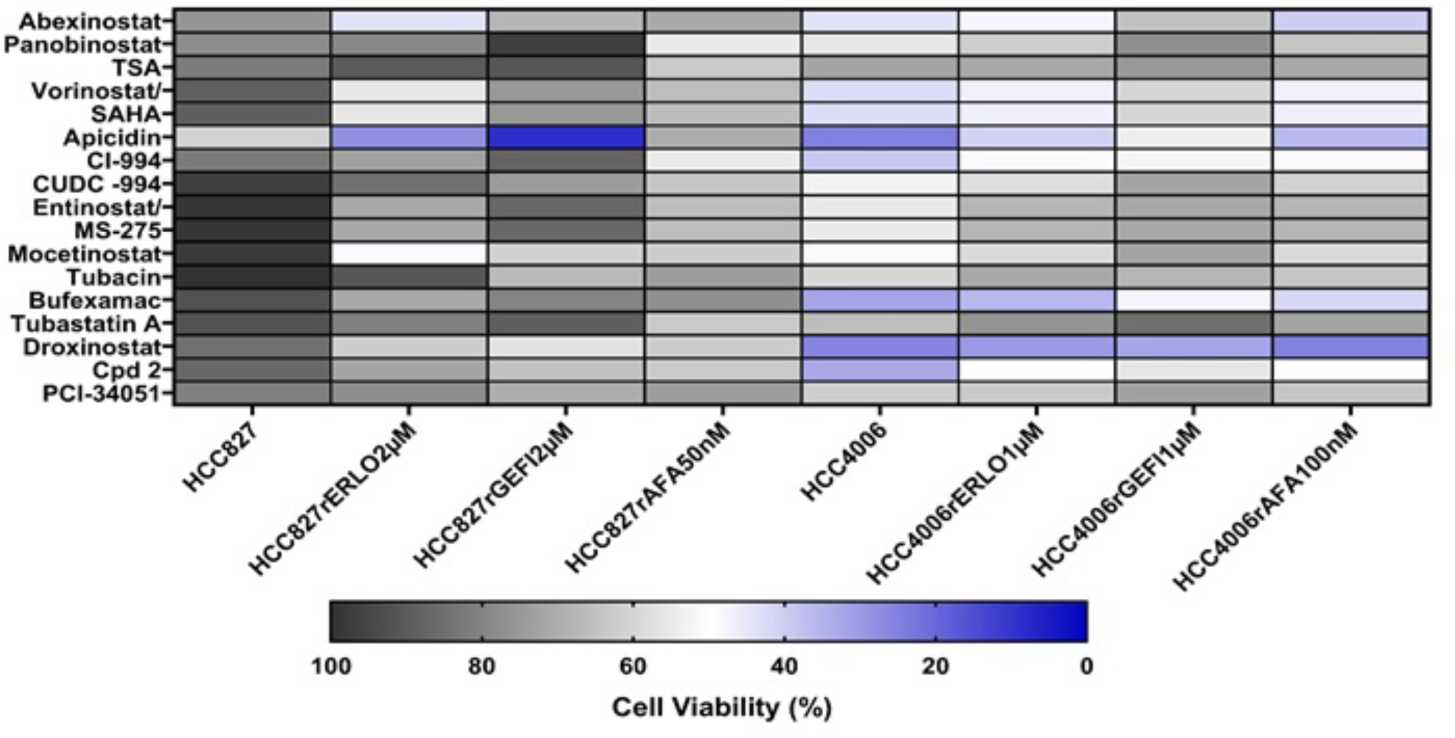
Effects of fixed HDAC inhibitor concentrations on cell viability. Cell viability was determined MTT assay after 120h of incubation. The rationale behind selecting the indicated concentrations is presented in Suppl. Figure 7. The numerical data are presented in Suppl. Table 8.

### Role of the individual cell line background in EGFR kinase resistance formation

The determination of drug response profiles indicated complex, individual phenotypes among the EGFR kinase inhibitor sublines. The correlation of the drug response profiles among the sublines adapted to the same drug demonstrated significant but not very pronounced correlations among the erlotinib- and gefitinib-resistant sublines and no significant correlation among the afatinib-adapted sublines (Figure 6A). The correlation of the drug response profiles among the sublines of the same parental cell lines demonstrated significant correlations among the HCC4006 sublines but not among the HCC827 sublines (Figure 6B). Based on these findings, it remains unclear if and if yes, to what extent, there are similarities between sublines adapted to the same drug and to what extent the cellular backgrounds contribute to the resistance phenotypes.

**Figure 6.**
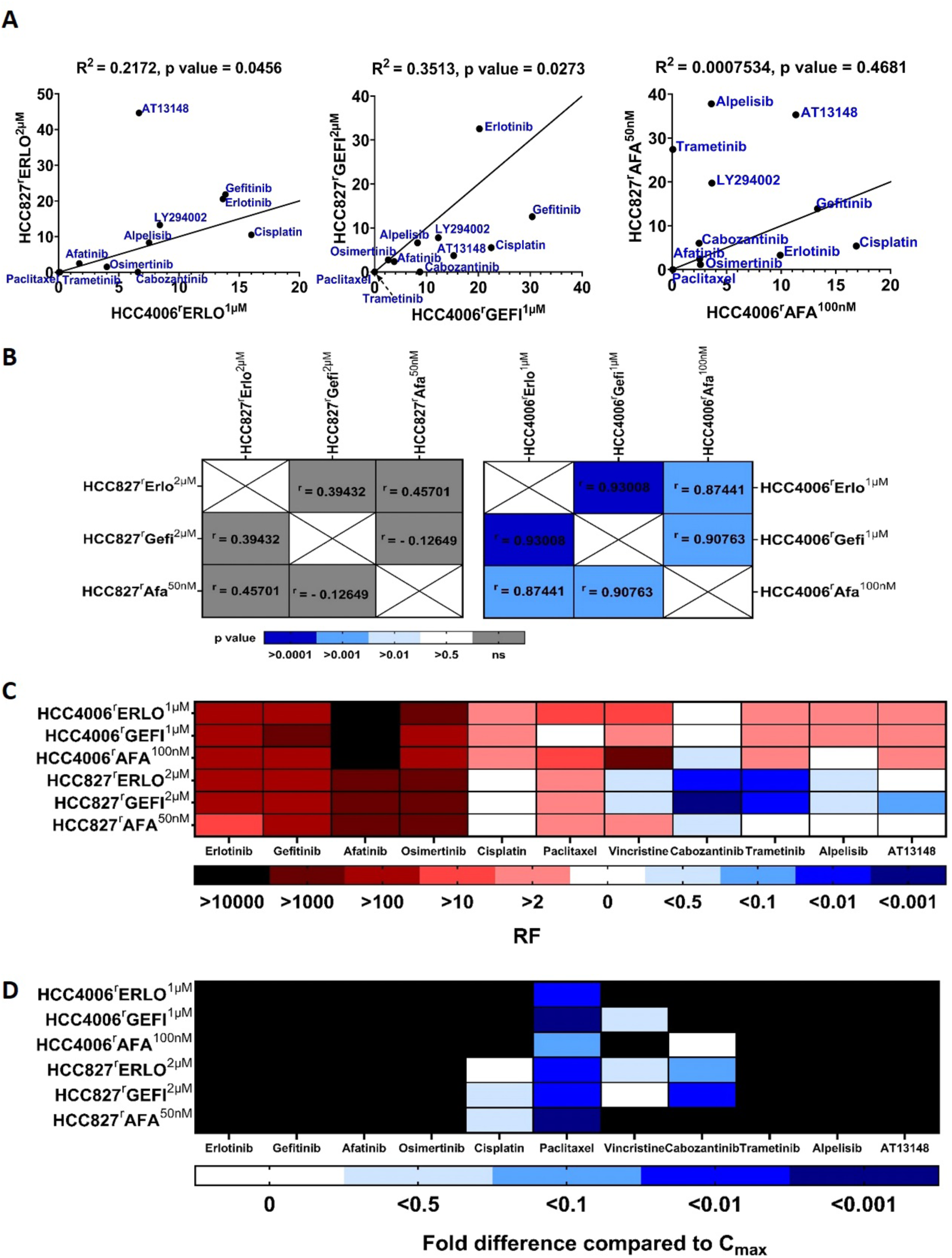
Correlation of the drug response patterns among the EGFR tyrosine kinase-adapted sublines. A) Correlation of the drug response profiles among the sublines adapted to the same drug. The straight line represents the best fit. B) Correlation of the drug response profiles among the sublines of the same parental cell lines. C) The heatmap summarising the drug sensitivity profiles of the EGFR tyrosine kinase inhibitor-adapted sublines relative to the respective parental cell lines expressed as resistance factors (RF). D) Heatmap summarising cell line sensitivity to therapeutic plasma concentrations (C_max_) of the indicated drugs. Black indicates IC_50_ values higher than the C_max_.

### Cross-resistance profiles in the context of clinically achievable therapeutic concentrations

The complexity of the phenotypes of the EGFR kinase inhibitor-adapted sublines can also be illustrated by an overarching heatmap reflecting the response to EGFR kinase inhibitors, additional kinase inhibitors, and cytotoxic anti-cancer drugs (Figure 6C). Moreover, when we considered the drug activities in the context of clinically achievable plasma concentrations (C_max_) (19), this interestingly resulted in different sensitivity patterns (Figure 6D). This indicates that drug response correlations derived from model systems are primarily insightful with regard to mechanistic considerations. For translational approaches, the clinically achievable plasma concentrations also need to be taken into account.

### Reversible shift in cell line oxygen consumption

Cancer cells may undergo a metabolic shift resulting in ATP production under normoxic conditions via glycolysis (instead of oxidative phosphorylation in the mitochondria), a phenomenon referred to as ‘aerobic glycolysis’ and/ or ‘Warburg effect’ (46). Such metabolic changes can also be associated with reduced cancer cell sensitivity to anti-cancer drugs, including EGFR tyrosine kinase inhibitors (46,47).

The MTT assay was used for the determination of drug effects in this study, which measures oxidative phosphorylation in the mitochondria as a surrogate for cell viability (34,48,49). Hence, the MTT assay is not suited for cells displaying a Warburg metabolism. During the course of the project, we occasionally observed that the signals from the cell untreated controls in HCC4006 and HCC827^r^GEFI^2μM^ did not significantly differ from the background signals of the cell culture media only controls, although microscopic inspection revealed viable, confluent cell layers in the respective former.

To investigate this phenomenon, we determined cellular oxygen consumption in the project cell lines. Moreover, the sensitivity of the project cell lines was tested against 2-Deoxy-D-Glucose (2DG), a competitive inhibitor of glycolysis with activity against cancer cells displaying a Warburg phenotype (50).

A standard respiration profile is provided in Figure 7A. Cells are monitored until a stable routine respiration level has been established. In cells with intact oxidative phosphorylation, the addition of the ATP synthase inhibitor oligomycin then causes a reduction in oxygen consumption (leak state). Next, the addition of FCCP (carbonyl cyanide 4-(trifluoromethoxy)phenylhydrazone), a protonophore that causes mitochondrial membrane permeabilisation, results in maximum respiration/ oxygen consumption (uncoupling, ETS = electron transfer system capacity). Finally, the complex III inhibitor antimycin A is added, resulting in the complete suppression of respiration, indicating non-mitochondrial oxygen consumption (Figure 7A). Small or no differences in oxygen consumption during the different assay steps indicate a lack of oxidative phosphorylation in the mitochondria.

**Figure 7.**
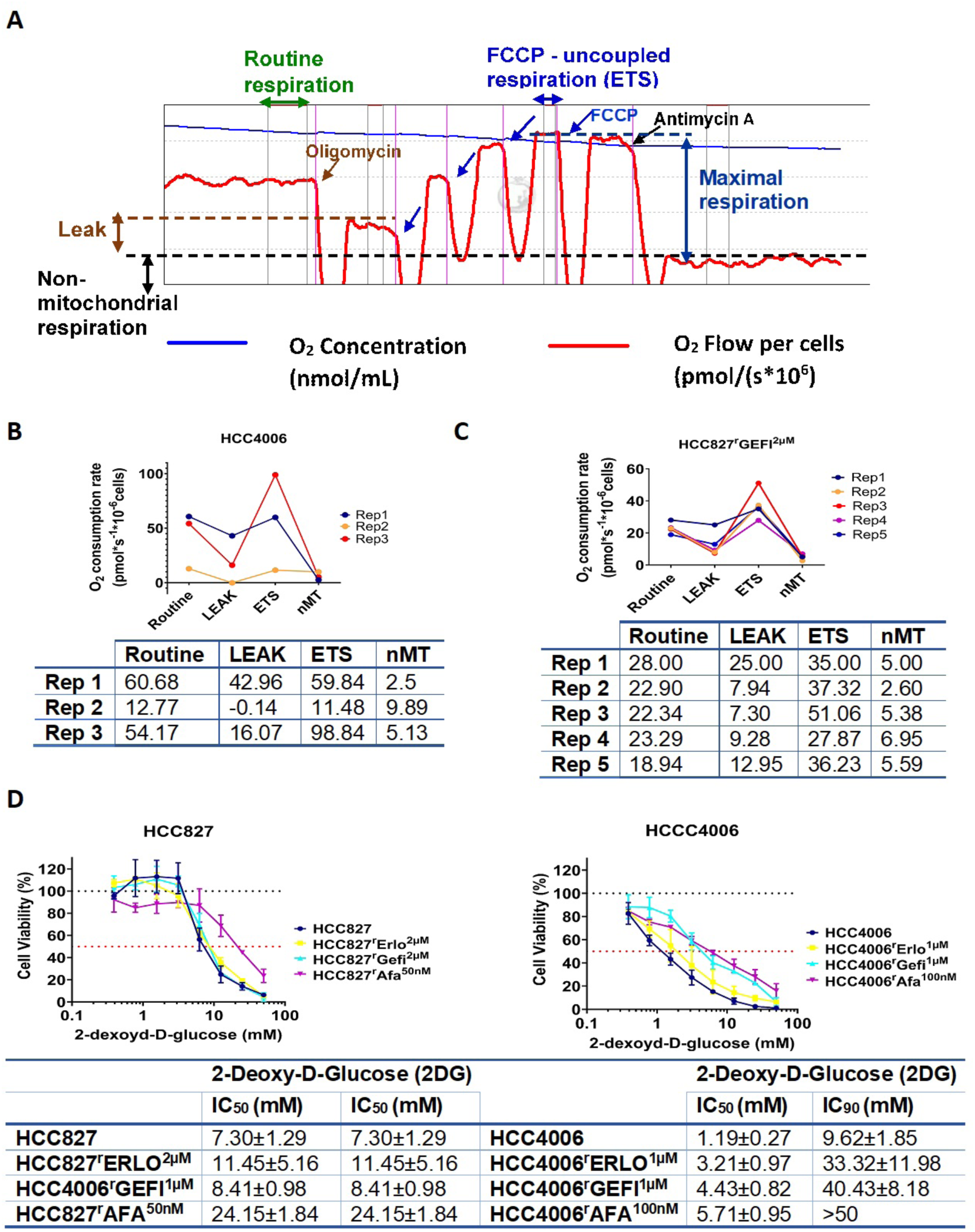
Measurement of oxygen consumption. Oxygen consumption was measured using an Oxygraph-2k respirometer. A) Representative high-resolution respirometry (HRR) profile of HCC827. Routine respiration is shown in green. Addition of the ATP synthase inhibitor oligomycin then causes a reduction in oxygen consumption (LEAK). FCCP (carbonyl cyanide 4-(trifluoromethoxy)phenylhydrazone) causes mitochondrial membrane permeabilization, resulting in maximum respiration/ oxygen consumption (uncoupling, ETS = electron transfer system capacity). Complex III inhibitor antimycin A addition result in respiration suppression, indicating non-mitochondrial oxygen consumption. B) Respirometry results of HCC4006 cells from three independent experiments. C) Respirometry results of HCC827^r^GEFI^2µM^ cells from three independent experiments. D) Effect of 2-Deoxy-D-Glucose (2DG) on cell line viability as indicated by MTT assay after 120h incubation. IC_50_ values were calculated using the software Calcusyn (Version 1.1, Biosof 1996).

Indeed, HCC4006 (Figure 7B) and HCC827^r^GEFI^2µM^ (Figure 7C) displayed temporary shifts towards a Warburg metabolism. However, the changes in metabolism occurred only sporadically and were unpredictable, which made it unfeasible to investigate this phenomenon systematically during this project. Nevertheless, the metabolism of these two cell lines seemed to fluctuate occasionally between oxidative phosphorylation and aerobic glycolysis. None of the other cell lines displayed similar shifts in their metabolism (Suppl. Figure 6).

2DG treatment resulted in complex results. HCC4006 and its sublines were generally more sensitive to HCC827 and its sublines, and HCC4006 was indeed the most 2DG-sensitive cell line of the panel (Figure 7D). However, the 2DG sensitivity of HCC827^r^GEFI^2µM^ was substantially lower, and similar to that of HCC827 (Figure 7D). Notably, we were not able to measure the 2DG sensitivity of HCC4006 and HCC827^r^GEFI^2µM^ when they were in an acute aerobic glycolysis metabolic phase. Further research will have to elucidate this metabolic plasticity of these cell lines further.

## Discussion

Here, we introduce a novel panel of NSCLC cell lines consisting of the EGFR-mutant cell lines HCC827 and HCC4006 and their sublines adapted to the EGFR tyrosine kinase inhibitors gefitinib (HCC827^r^GEFI^2µM^, HCC4006^r^GEFI^1µM^), erlotinib (HCC827^r^ERLO^2µM^, HCC4006^r^ERLO^1µM^), and afatinib (HCC827^r^AFA^50nM^, HCC4006^r^AFA^100nM^). Notably, all resistant sublines displayed resistance to gefitinib, erlotinib, and afatinib and also to the third-generation EGFR kinase inhibitor osimertinib, which was developed to overcome resistance mediated by T790M EGFR mutations (2,5). Hence, these findings indicate that the resistance in the EGFR kinase inhibitor-adapted sublines is not mediated by T790M mutations, but rather by EGFR-independent mechanisms that mediate resistance to first, second, and third generation EGFR tyrosine kinase inhibitors (2,5,51). Hence, our results also indicate that while acquired EGFR kinase inhibitor resistance may be delayed in some cases by designing further generations of EGFR kinase inhibitors that target additional EGFR resistance mutations (51), resistance is likely to emerge eventually by EGFR-independent processes. Accordingly, the combination of amivantamab, a bispecific antibody targeting EGFR and MET, with the third generation EGFR kinase inhibitor lazertinib was found superior to lazertinib or osimertinib alone (52).

Resistance formation resulted in morphological changes in some of the sublines. Most notably, HCC4006^r^ERLO^1µM^ displayed a more spindle-like morphology compared to its parental cell line HCC4006. This is in agreement with findings that had suggested that the HCC4006^r^ERLO^1µM^ precursor cell line HCC4006^r^ERLO^0.5µM^ displayed an epithelial-mesenchymal-transition (EMT) phenotype (53). However, the same study had also reported EMT for the HCC4006^r^GEFI^1µM^ precursor cell line HCC4006^r^GEFI^0.5µM^ and for HCC4006^r^AFA^100nM^ (53), but we did not find any morphological indications of this in HCC4006^r^GEFI^1µM^ or HCC4006^r^AFA^100nM^. This suggests that EMT can be dynamically regulated during the continued drug adaptation process and that there may be some level of plasticity even in the established resistant cell lines. Such plasticity may in addition to different laboratory routines and cell line evolution contribute to differing phenotypes among identical cell lines, as previously shown in HeLa and other cancer cell lines (54,55).

Previous studies had further reported that the HCC4006^r^ERLO^1µM^ precursor cell line HCC4006^r^ERLO^0.5µM^ was characterised by Shc-initated RAS/RAF/MEK/ERK signalling and CIP2A-mediated AKT activation and that MEK and AKT inhibitors inhibited the growth of this cell line in a similar way as the parental HCC4006 cell line (56,57). In contrast, HCC4006^r^ERLO^1µM^ displayed increased resistance to the MEK inhibitor trametinib relative to HCC4006, suggesting that survival of this cell line is not driven by MEK signalling. Moreover, while HCC4006^r^ERLO^1µM^ displayed similar sensitivity to the PI3K inhibitor LY294002 as HCC4006, it was cross-resistant to the PI3K inhibitor alpelisib relative to HCC4006. Again, these discrepancies between HCC4006^r^ERLO^0.5µM^ and HCC4006^r^ERLO^1µM^ may indicate a pronounced level of cancer cell plasticity during the resistance formation process.

We further detected metabolic plasticity in HCC4006 and HCC827^r^GEFI^2µM^ during our project. Both cell lines temporarily displayed a Warburg metabolism, i.e. they did not consume oxygen despite its presence. Notably, we have previously shown that drug-response data for individual drug/ cell line combinations from the NCI60 screen was characterised by a very high level of variability (58). Cancer cell line plasticity resulting in phenotypic changes during cultivation may contribute to these variations.

Generally, the determination of response profiles to cytotoxic anti-cancer drugs, kinase inhibitors, and HDAC inhibitors resulted in complex patterns that were specific for each individual subline without obvious overlaps. This suggests that each resistance formation process follows its own unpredictable route. Notably, these findings are not only in line with other studies investigating drug-resistant cancer cell lines, including those, in which the same cell line was repeatedly adapted to the same drug in multiple experiments (18,59–64), but also with the complex evolutionary processes in cancer cells from lung cancer patients (65–69).

As a side aspect, we observed that drug response patterns differed, when we did not directly correlate the drug effects but considered drug efficacy in the context of the respective clinically achievable (therapeutic) drug concentrations. Hence, direct drug response correlations may be of mechanistic relevance, but therapeutic concentrations need to be considered for translational approaches.

Notably, all resistant sublines remained sensitive or even displayed collateral sensitivity to at least one of the investigated drugs. This suggests that there are, in principle, effective treatments available for cancer cells that have acquired resistance to a certain drug. Further research will have to develop an improved understanding of the determinants of drug sensitivity in cancer cells that have stopped responding to the normally applied standard therapies, including biomarkers that guide effective next-line therapies to patients, who are likely to benefit from them.

In conclusion, we here introduce a novel panel of NSCLC cell lines with acquired resistance to EGFR kinase inhibitors. Drug response profiles differed significantly between the resistant sublines, suggesting that each resistance formation process follows an individual, unpredictable path. The comparison of some of the sublines with precursor cell lines that had been previously characterised at a lower resistance level indicated a substantial level of phenotypic heterogeneity during the ongoing resistance formation process. Moreover, HCC4006 and HCC827^r^GEFI^2µM^ displayed metabolic plasticity during the course of our experiments. This suggests that cancer cells are subject to a continuous plasticity that affects their drug sensitivity profiles and may contribute to the variability in cell line phenotypes observed between different laboratories and also in intra-laboratory experiments (54,55,58). Future research will have to establish a detailed understanding of these dynamic processes that can be translated into biomarker-guided strategies that guide effective therapies to patients with therapy-refractory disease for whom currently no standard treatment options are available.

## Materials and Methods

### Cell culture

HCC4006 was purchased from ATCC (Manassas, VA, USA), HCC827 from DSMZ (Braunschweig, Germany). The drug-adapted sub-lines were established by continuous exposure to stepwise increasing drug concentrations as previously described (59) and derived from the Resistant Cancer Cell Line Collection (RCCL), (https://research.kent.ac.uk/industrial-biotechnology-centre/the-resistant-cancer-cell-line-rccl-collection/).

All cell lines were cultured in Iscove’s Modified Dulbecco’s Medium (IMDM; Gibco^TM^, Life technologies, UK), supplemented with 10% (v/v) foetal bovine serum (FBS; Sigma-Aldrich, UK), and 100 IU/mL penicillin and 100µg/mL of streptomycin (Gibco^TM^, Life technologies, UK), at 37°C in a humidified 5% CO_2_ incubator. The media for resistant cell lines was additionally supplemented with the respective adaptation drug concentrations as specified by the cell line name. For example, HCC4006^r^AFA^100nM^ was maintained in 100nM afatinib.

### Compounds

The following compounds were obtained from specified suppliers: Erlotinib, gefitinib, Afatinib, Osimertinib, Cisplatin, Paclitaxel (Selleckchem), Cabozantinib, Trametinib, Alpelisib, LY294002, Zosuquidar (MedChemExpress), AT13148 (Astex Pharmaceuticals), Vincristine (Cayman Chemicals), 2-Deoxy-D-Glucose (Sigma-Aldrich). Most of the compounds were prepared and diluted using dimethyl sulfoxide (DMSO; Sigma-Aldrich, Germany) under sterile conditions and stored at -80°C, except for cisplatin which was dissolved in 0.9% saline solution (0.9% (w/v) and stored in dark tubes at room temperature.

### Cell imaging

The cell images were captured using bright field microscopy with a GXCAM-U3-5 industrial camera using different magnifications (Olympus CKX53 inverted microscope, Olympus Life Sciences, UK).

### Growth kinetics

The xCelligence real time cell analyser (RTCA) system was used to investigate the growth kinetics according to the manufacturer’s instructions. Cells were grown in 16-well microtiter plates (E-Plate, ACEA Biosciences Inc. San Diego, CA, USA) in three replicates.

### Cell viability assays

If not stated differently, cell viability was determined by (3-(4,5-dimethylthiazol-2-yl)-2,5-diphenyltetrazolium bromide) MTT assay after 120-hour incubation as previously described (34). Alternatively, the sulforhodamine B (SRB) assay performed. Cells were fixed using 10% (w/v) trichloroacetic acid (TCA) and stained with 0.4% (w/v) SRB. Protein-bound to SRB was then solubilised by adding 100µL 10mM Tris-base per well (ThermoFisher Scientific, USA). Absorbance was determined using a Victor X4 Multilabel plate reader (PerkinElmerLife Sciences, UK) at 490nM. The percentage viability of drug treated cells was calculated relative to the untreated control. Then, the half maximal inhibitory concentrations (IC_50_) of the specific drug were determined using CalcuSyn software (Biosoft, UK).

### Determination of oxygen consumption rates

Oxygen consumption in cells was determined using Oxygraph-2k by Oroboros (Oroboros O2k, Oroboros Instruments GmbH, Austria) following the manufacturer’s instructions. 1 × 10^6^ cells per mL cell suspensions were used per experiment. Oxygen consumption was monitored using the O2k software DatLab. After the system had reached an equilibrium (routine/basis respiration value), 2µL of a (4mg/mL) solution of the ATP synthase inhibitor oligomycin (Sigma Aldrich, Germany) was added to determine the oligomycin-insensitive reduction in oxygen consumption (leak state). Then, 2µL of 200µM carbonyl cyanide 4-(trifluoromethoxy)phenylhydrazone (FCCP) (Sigma Aldrich, Germany) solution was added to determine the maximum mitochondrial uncoupled respiration rate, referred to as the electron transport system value (ETS). Finally, 2µL of 2mM antimycin A (Sigma Aldrich, Germany) was added to measure the non-mitochondrial oxygen consumption values (nMT). From the results, the ratios of mitochondrial routine respiration to ETS (Routine:ETS) and leak respiration to ETS (Leak:ETS) were calculated.

### Statistical analysis and data manipulation

Statistical analysis was performed using GraphPad Prism 10 (GraphPad Software, Inc., USA). Two-tailed t-test used for single comparison assuming unequal variance. Multiple comparison was analysed using two-way analysis of variance (ANOVA) followed by post-hoc testing by Tukey’s pairwise comparison/ Dunnett’s multiple comparison with 95% confidence interval.

## Availability of data and materials

All data are provided in the manuscript and the supplements.

## Competing interests

Nothing to declare.

## Funding

This work was supported by grants from the Frankfurter Stiftung für krebskranke Kinder.

## Acknowledgements

We would like to thank Astex Pharmaceuticals for their kind gift of AT13148.

## Notes

### Competing Interest Statement

The authors have declared no competing interest.

