## Supplementary Figures and Tables for "Phenotypic plasticity in a novel set of EGFR tyrosine kinase inhibitor-adapted non-small cell lung cancer cell lines"

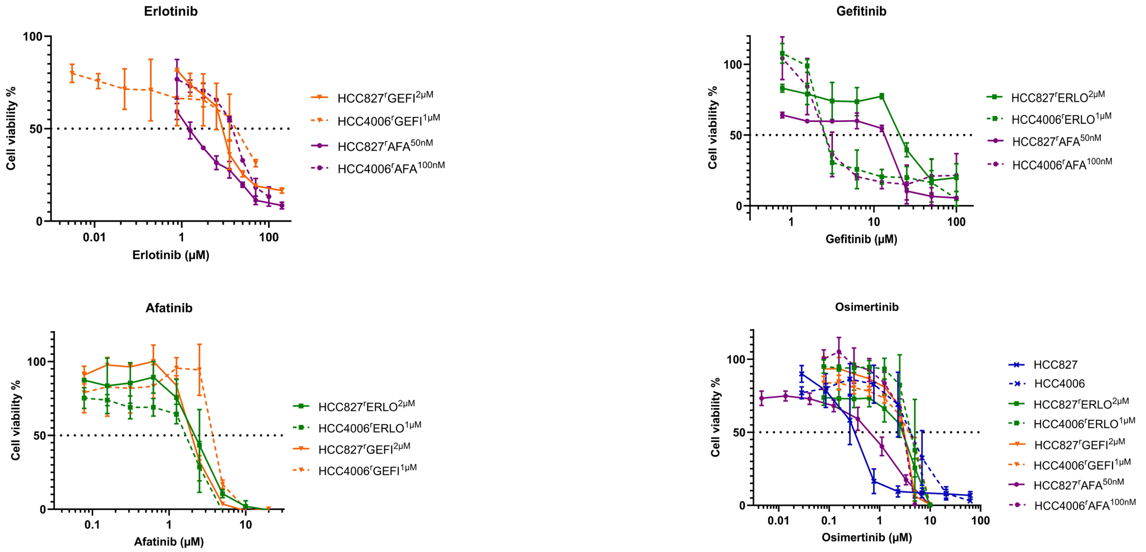


**Supplementary Figure 1. Dose response curves of HCC827 and HCC4006 and their EGFR tyrosine kinase inhibitor-resistant sublines to different EGFR tyrosine kinase inhibitors.** Data points represent means of at least three biological repeats ± S.D, as determined by MTT assay after a 120h incubation period.

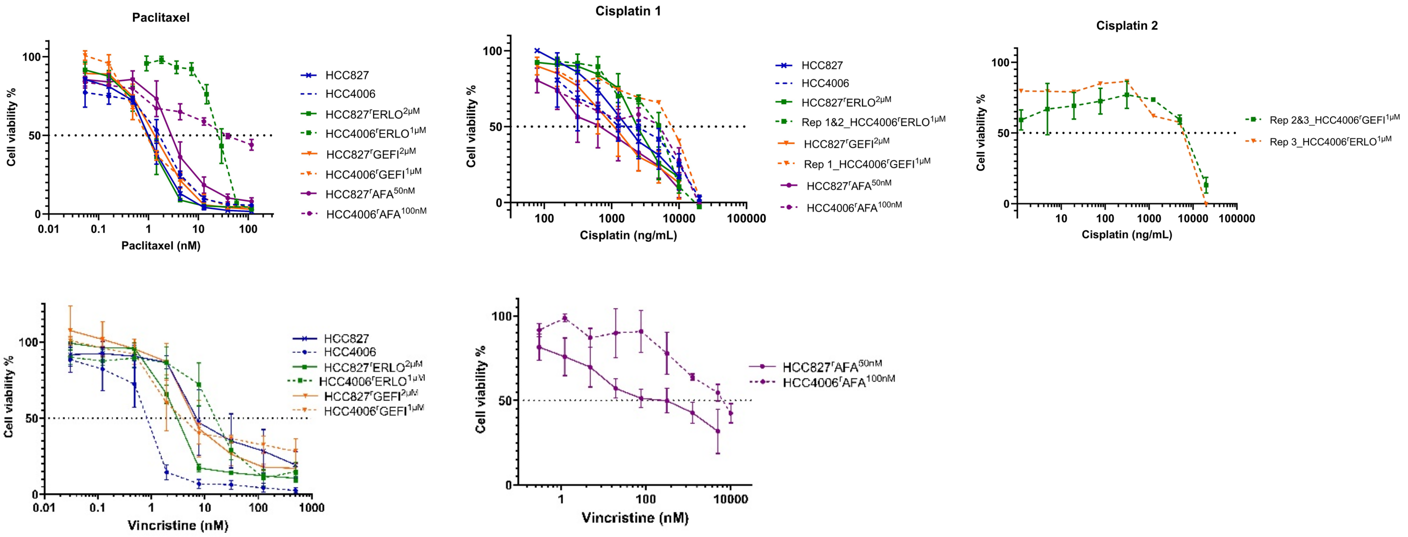


**Supplementary Figure 2. Dose response curves of HCC827 and HCC4006 and their EGFR tyrosine kinase inhibitor-resistant sublines to different cytotoxic anti-cancer drugs.** Data points represent means of at least three biological repeats ± S.D, as determined by MTT assay after a 120h incubation period.

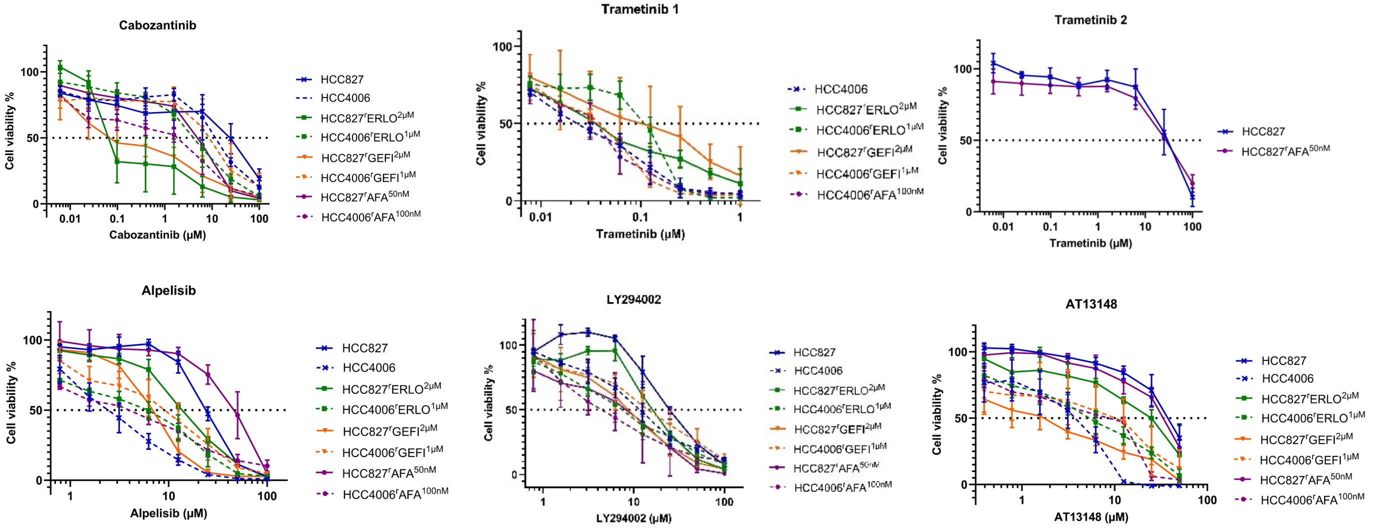


**Supplementary Figure 3. Dose response curves of HCC827 and HCC4006 and their EGFR tyrosine kinase inhibitor-resistant sublines to different kinase inhibitors.** Data points represent means of at least three biological repeats ± S.D, as determined by MTT assay after a 120h incubation period.

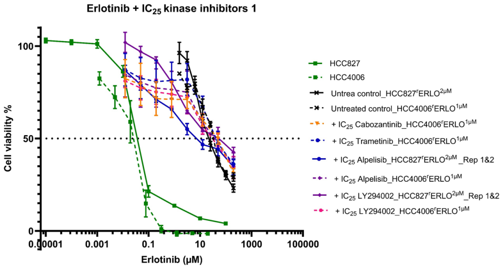
Erlotinib + IC25 kinase inhibitors 1
Cell viability %
Erlotinib (µM)
HCC827
HCC4006
Untrea control_HCC827rERLO2µM
Untreated control_HCC4006rERLO1µM
+ IC25 Cabozantinib_HCC4006rERLO1µM
+ IC25 Trametinib_HCC4006rERLO1µM
+ IC25 Alpelisib_HCC827rERLO2µM_Rep 1&2
+ IC25 Alpelisib_HCC4006rERLO1µM
+ IC25 LY294002_HCC827rERLO2µM_Rep 1&2
+ IC25 LY294002_HCC4006rERLO1µM

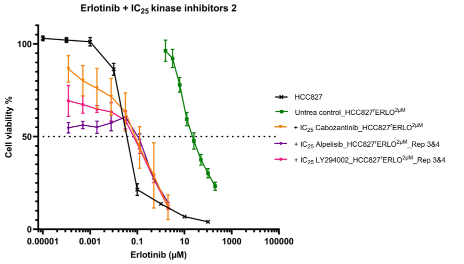
Erlotinib + IC25 kinase inhibitors 2
Cell viability %
Erlotinib (µM)
HCC827
Untrea control_HCC827rERLO2µM
+ IC25 Cabozantinib_HCC827rERLO2µM
+ IC25 Alpelisib_HCC827rERLO2µM_Rep 3&4
+ IC25 LY294002_HCC827rERLO2µM_Rep 3&4

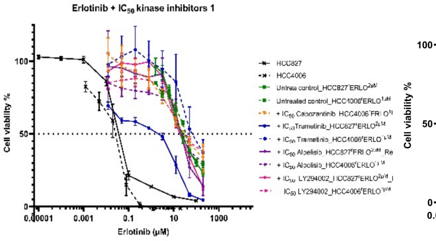
Erlotinib + IC50 kinase inhibitors 1
Cell viability %
Erlotinib (µM)
HCC827
HCC4006

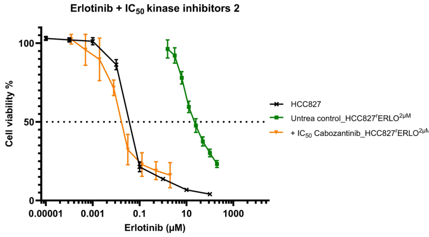
Erlotinib + IC50 kinase inhibitors 2
Cell viability %
Erlotinib (µM)
HCC827
Untrea control_HCC827rERLO2µM

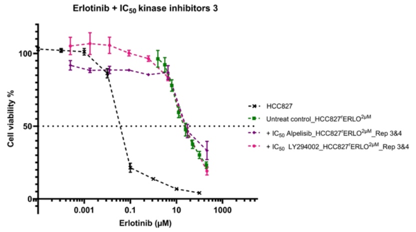
Erlotinib + IC50 kinase inhibitors 3
Cell viability %
Erlotinib (µM)
HCC4006
Untreat control_HCC827rERLO2µM
+ IC50 Alpelisib_HCC827rERLO2µM_Rep 3&4
+ IC50 LY294002_HCC827rERLO2µM_Rep 3&4

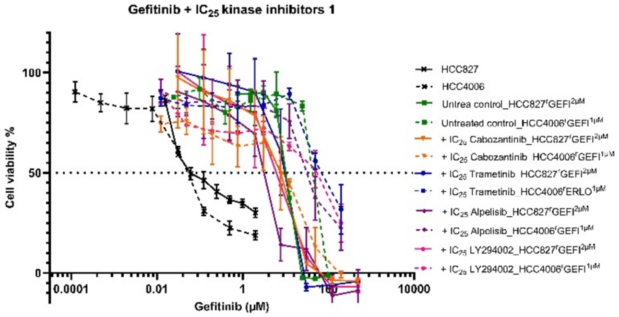
Gefitinib + IC25 kinase inhibitors 1
Cell viability %
Gefitinib (µM)
HCC827
HCC4006
Untrea control_HCC827rGEFI2µM
Untreated control_HCC4006rGEFI1µM
+ IC25 Cabozantinib_HCC827rGEFI2µM
+ IC25 Trametinib_HCC827rGEFI2µM
+ IC25 Trametinib_HCC4006rGEFI1µM
+ IC25 Alpelisib_HCC827rGEFI2µM
+ IC25 Alpelisib_HCC4006rGEFI1µM
+ IC25 LY294002_HCC827rGEFI2µM
+ IC25 LY294002_HCC4006rGEFI1µM

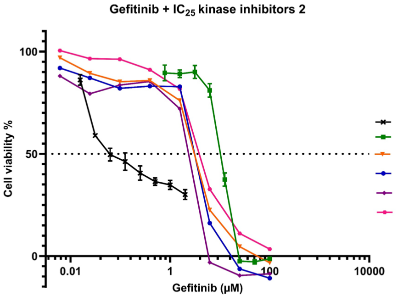
Gefitinib + IC25 kinase inhibitors 2
Cell viability %
Gefitinib (µM)
HCC827
Untrea control_HCC827rGEFI2µM
+ IC25 Cabozantinib_HCC827rGEFI2µM_Rep 3
+ IC25 Trametinib_HCC827rGEFI2µM_Rep 3
+ IC25 Alpelisib_HCC827rGEFI2µM_Rep 3
+ IC25 LY294002_HCC827rGEFI2µM_Rep 3

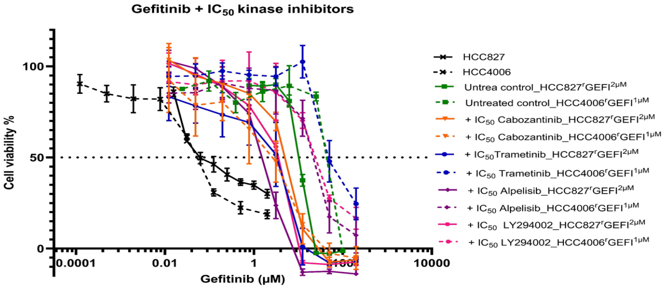
Gefitinib + IC50 kinase inhibitors
Cell viability %
Gefitinib (µM)
HCC827
HCC4006
Untrea control_HCC827rGEFI2µM
Untreated control_HCC4006rGEFI1µM
+ IC50 Cabozantinib_HCC827rGEFI2µM
+ IC50 Cabozantinib_HCC4006rGEFI1µM
+ IC50Trametinib_HCC827rGEFI2µM
+ IC50 Trametinib_HCC4006rGEFI1µM
+ IC50 Alpelisib_HCC827rGEFI2µM
+ IC50 Alpelisib_HCC4006rGEFI1µM
+ IC50 LY294002_HCC827rGEFI2µM
+ IC50 LY294002_HCC4006rGEFI1µM

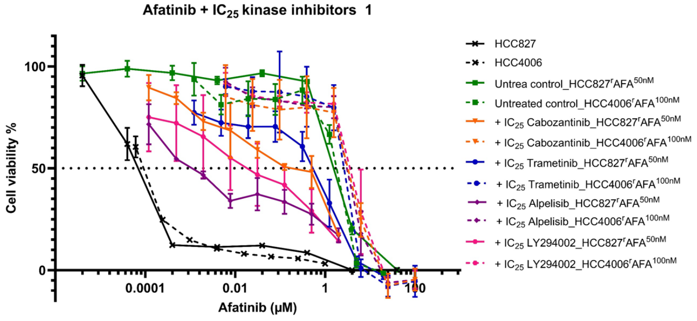
Afatinib + IC25 kinase inhibitors 1
Cell viability %
Afatinib (µM)
HCC827
HCC4006
Untrea control_HCC827rAFA50nM
Untreated control_HCC4006rAFA100nM
+ IC25 Cabozantinib_HCC827rAFA50nM
+ IC25 Cabozantinib_HCC4006rAFA100nM
+ IC25 Trametinib_HCC827rAFA50nM
+ IC25 Trametinib_HCC4006rAFA100nM
+ IC25 Alpelisib_HCC827rAFA50nM
+ IC25 Alpelisib_HCC4006rAFA100nM
+ IC25 LY294002_HCC827rAFA50nM
+ IC25 LY294002_HCC4006rAFA100nM

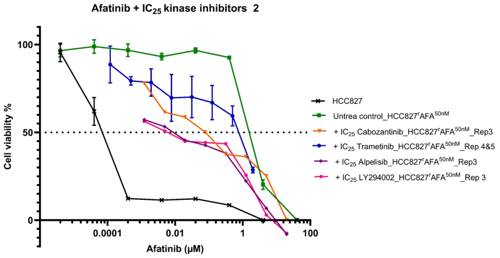
Afatinib + IC25 kinase inhibitors 2
Cell viability %
Afatinib (µM)
HCC827
Untrea control_HCC827rAFA50nM
+ IC25 Cabozantinib_HCC827rAFA50nM_Rep3
+ IC25 Trametinib_HCC827rAFA50nM_Rep 4&5
+ IC25 Alpelisib_HCC827rAFA50nM_Rep3
+ IC25 LY294002_HCC827rAFA50nM_Rep 3

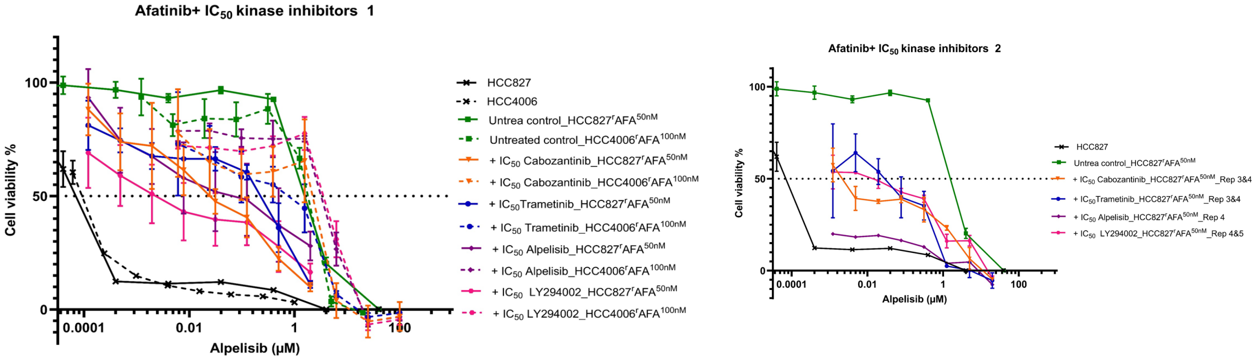


**Supplementary Figure 4. Effects of kinase inhibitors on the sensitivity of EGFR tyrosine kinase-adapted sublines to the respective drugs of adaptation.** Data points represent means of at least three biological repeats ± S.D, as determined by MTT assay after a 120h incubation period.

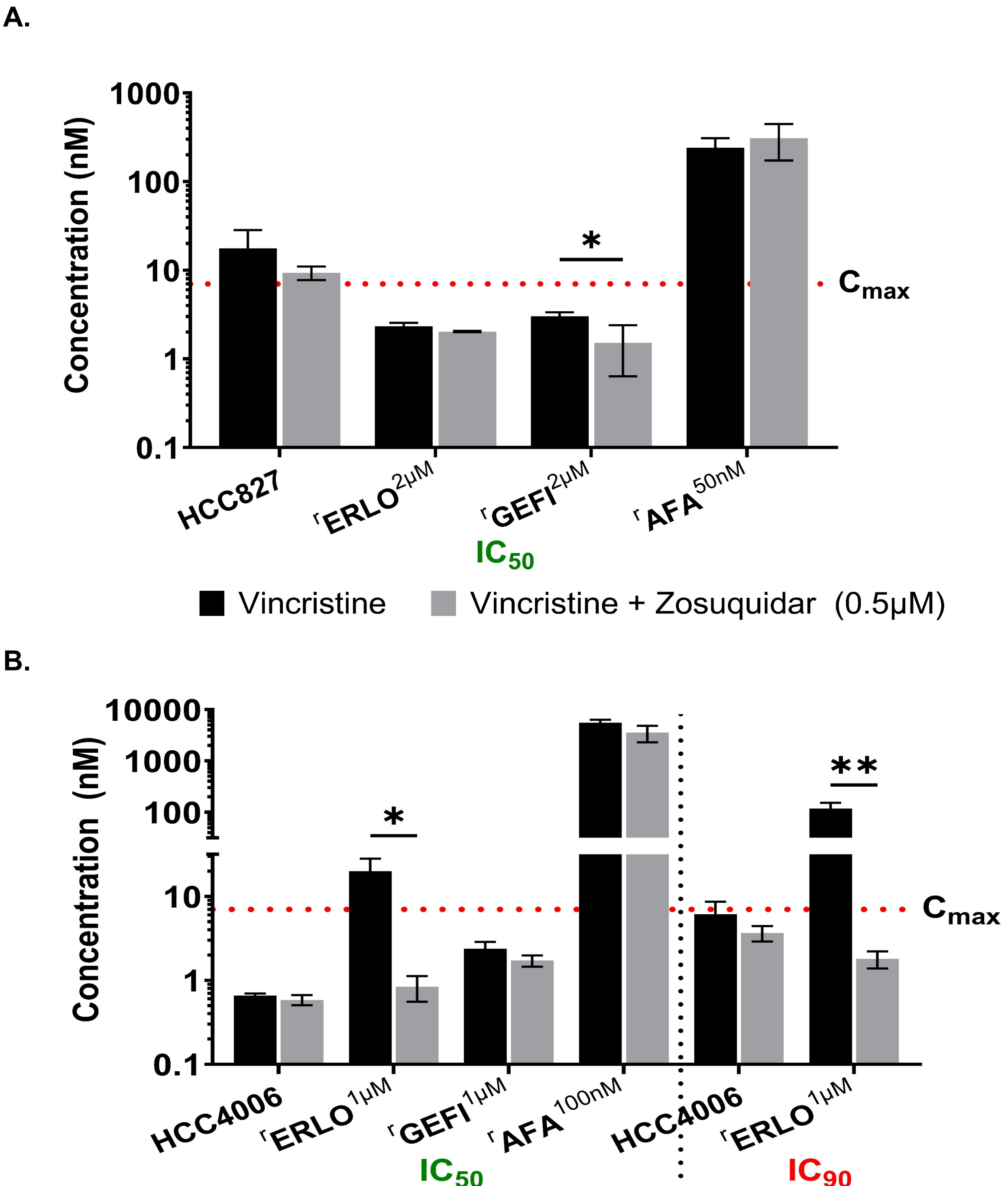


**Supplementary Figure 5. Determination of $IC_{50}$ and $IC_{90}$ value of vincristine in HCC4006 and HCC827 and their EGFR tyrosine kinase-adapted sublines in the presence or absence of the ABCB1 inhibitor zosuquidar.** Values represent means of at least three biological repeats ± S.D. Drug response was determined by MTT assay after a 120h incubation period. $IC_{50}$ values were calculated using Calcusyn (Version 1.1, Biosof 1996).

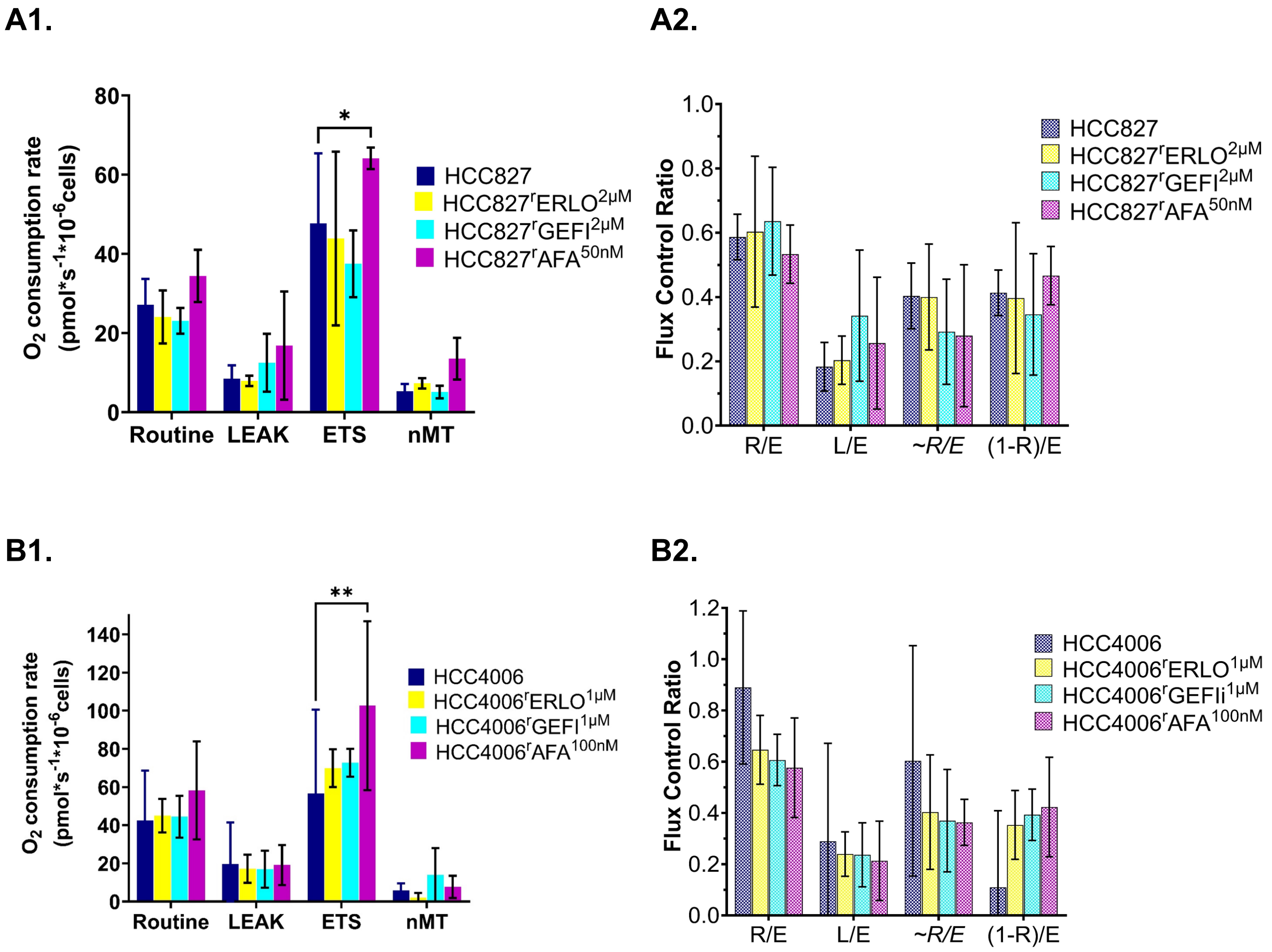


**Supplementary Figure 6. Summary of high-resolution respirometry results of HCC4006 and its EGFR tyrosine kinase inhibitor-adapted sublines.** Data are presented as oxygen consumption rates and flux control ratios. Bars represent the mean ± S.D. of three independent experiments.

**Supplementary Table 1. Doubling times of HCC827 and HCC4006 and their EGFR tyrosine kinase inhibitor-adapted sublines in the absence and presence of drug.** Values represent the means of at least three biological replicates ± S.D. Statistical significance was tested using two-way ANOVA with post-hoc Tukey's pairwise comparison test. * $p<0.05$

| | Doubling time (Hrs) | | | |
|---|---|---|---|---|
| | HCC827 | HCC827$^{r}$ERLO$^{2\mu M}$ | HCC827$^{r}$GEFI$^{2\mu M}$ | HCC827$^{r}$AFA$^{50nM}$ |
| No drug | 12.8 ± 0.8 | 18.6 ± 4.3 | 20.2 ± 7.3 | 15.1 ± 5.6 |
| With drug | - | 23.4 ± 2.8* | 28.8 ± 0.5* | 20.9 ± 4.4 |

| | Doubling time (Hrs) | | | |
|---|---|---|---|---|
| | HCC4006 | HCC4006$^{r}$ERLO$^{1\mu M}$ | HCC4006$^{r}$GEFIi$^{1\mu M}$ | HCC4006$^{r}$AFA$^{100nM}$ |
| No drug | 25.4 ± 3.4 | 28.3 ± 6.4 | 23.2 ± 3.0 | 31.5 ± 7.4 |
| With drug | n/a | 29.7 ± 4.8 | 27.8 ± 11.9 | 35.0 ± 16.4 |

**Supplementary Table 2. Sensitivity of HCC827, HCC4006, and their EGFR tyrosine kinase inhibitor-resistant sublines to EGFR tyrosine kinase inhibitors.** $IC_{50}$ values represent means of at least three biological repeats ± S.D. Drug response was determined by MTT assay after a 120h incubation period. $IC_{50}$ values were calculated using Calcusyn (Version 1.1, Biosof 1996).

| Cell line | Drug | | | | | | | |
|---|---|---|---|---|---|---|---|---|
| | **Erlotinib** | | **Gefitinib** | | **Afatinib** | | **Osimertinib** | |
| | **$IC_{50}$** | **RF** | **$IC_{50}$** (µM) | **RF** | **$IC_{50}$** (nM) | **RF** | **$IC_{50}$** (µM) | **RF** |
| HCC827 | 0.08±0.02 | n/a | 0.08±0.02 | n/a | 1.32±0.47 | n/a | 0.0004±0.0002 | n/a |
| HCC827$^{r}$ERLO$^{2µM}$ | 22.02±3.19 | 282.6 | 21.80±1.45 | 290.5 | 2232.96±628.34 | 1690.7 | 1.53±0.36 | 3642.9 |
| HCC827$^{r}$GEFI$^{2µM}$ | 32.54±6.28 | 417.6 | 13.36±1.58 | 178.1 | 2359.51±238.97 | 1786.5 | 2.74±0.42 | 6523.8 |
| HCC827$^{r}$AFA$^{50nM}$ | 3.29±1.19 | 42.2 | 13.88±1.50 | 185.0 | 2417.76±262.63 | 1830.6 | 1.14±0.31 | 2714.3 |

| Cell line | Drug | | | | | | | |
|---|---|---|---|---|---|---|---|---|
| | **Erlotinib** | | **Gefitinib** | | **Afatinib** | | **Osimertinib** | |
| | **$IC_{50}$** (µM) | **RF** | **$IC_{50}$** (µM) | **RF** | **IC50** (nM) | **RF** | **$IC_{50}$** (µM) | **RF** |
| HCC4006 | 0.03±0.01 | n/a | 0.03±0.00 | n/a | 0.15±0.05 | n/a | 0.004±0.003 | n/a |
| HCC4006$^{r}$ERLO$^{1µM}$ | 13.65±4.19 | 519.7 | 13.86±1.59 | 473.1 | 1715.55±308.09 | 11319.0 | 4.01±1.35 | 1000.8 |
| HCC4006$^{r}$GEFI$^{1µM}$ | 20.20±2.52 | 769.3 | 33.38±14.03 | 1139.9 | 3761.06±883.09 | 24815.1 | 2.62±0.49 | 653.9 |
| HCC4006$^{r}$AFA$^{100nM}$ | 11.18±3.98 | 425.9 | 13.33±0.14 | 455.0 | 2533.50±664.04 | 16715.8 | 2.60±0.25 | 650.0 |

**Supplementary Table 3. Sensitivity of HCC827, HCC4006, and their EGFR tyrosine kinase inhibitor-resistant sublines to cytotoxic anti-cancer drugs.** $IC_{50}$ values represent means of at least three biological repeats ± S.D. Drug response was determined by MTT assay after a 120h incubation period. $IC_{50}$ values were calculated using Calcusyn (Version 1.1, Biosof 1996).

| Cell line | Drug | | | | | |
|---|---|---|---|---|---|---|
| | **Cisplatin** | | **Paclitaxel** | | **Vincristine** | |
| | **$IC_{50}$** (ng/mL) | **RF** | **$IC_{50}$** (nM) | **RF** | **$IC_{50}$** (nM) | **RF** |
| HCC827 | 2353.34±235.15 | n/a | 0.90±0.11 | n/a | 18.58±9.47 | n/a |
| HCC827$^{r}$ERLO$^{2μM}$ | 3166.98±596.87 | 1.35 | 5.58±0.91 | 6.20 | 2.96±0.40 | 0.16 |
| HCC827$^{r}$GEFI$^{2μM}$ | 1430.77±531.86 | 0.61 | 6.74±0.22 | 7.49 | 6.19±0.84 | 0.33 |
| HCC827$^{r}$AFA$^{50nM}$ | 1617.92±662.61 | 0.69 | 3.04±1.44 | 3.38 | 126.78±53.01 | 8.04 |

| Cell line | Drug | | | | | |
|---|---|---|---|---|---|---|
| | **Cisplatin** | | **Paclitaxel** | | **Vincristine** | |
| | **$IC_{50}$** (ng/mL) | **RF** | **$IC_{50}$** (nM) | **RF** | **$IC_{50}$** (nM) | **RF** |
| HCC4006 | 1477.36±540.32 | n/a | 1.14±0.20 | n/a | 0.84±0.21 | n/a |
| HCC4006$^{r}$ERLO$^{1μM}$ | 4819.52±853.62 | 3.26 | 24.71±4.18 | 21.68 | 14.83±9.16 | 17.55 |
| HCC4006$^{r}$GEFI$^{1μM}$ | 6768.33±854.69 | 4.58 | 1.09±0.03 | 0.96 | 1.74±0.67 | 2.06 |
| HCC4006$^{r}$AFA$^{100nM}$ | 5082.04±342.74 | 3.44 | 70.06±19.69 | 61.46 | 4604±120 | 5451 |

**Supplementary Table 4. Sensitivity of HCC827, HCC4006, and their EGFR tyrosine kinase inhibitor-resistant sublines to different kinase inhibitors.** $IC_{50}$ values represent means of at least three biological repeats ± S.D. Drug response was determined by MTT assay after a 120h incubation period. $IC_{50}$ values were calculated using Calcusyn (Version 1.1, Biosof 1996). Resistance factors (RF) were calculated as follows $IC_{50}$ drug-resistant subline/ $IC_{50}$ respective parental cell line.

| Cell line | Drug | | | | | | | | | |
|---|---|---|---|---|---|---|---|---|---|---|
| | Cabozantinib | | Trametinib | | Alpelisib | | LY294002 | | AT13148 | |
| | **$IC_{50}$** (μM) | **RF** | **$IC_{50}$** (μM) | **RF** | **$IC_{50}$** (μM) | **RF** | **$IC_{50}$** (μM) | **RF** | **$IC_{50}$**(μM) | **RF** |
| HCC827 | 33.99±9.19 | n/a | 25.56±2.88 | n/a | 28.73±0.98 | n/a | 26.10±1.40 | n/a | 43.89 | n/a |
| HCC827$^r$ERLO$^{2\mu M}$ | 0.062±0.032 | 0.0018 | 0.035 ±0.003 | 0.0014 | 9.45±2.43 | 0.33 | 13.54±0.82 | 0.52 | 44.64 | 1.02 |
| HCC827$^r$GEFI$^{2\mu M}$ | 0.009±0.003 | 0.0004 | 0.061±0.034 | 0.0024 | 6.82±0.67 | 0.24 | 7.79±1.75 | 0.30 | 3.71 | 0.08 |
| HCC827$^r$AFA$^{50nM}$ | 6.03±0.53 | 0.18 | 29.486±4.18 | 1.15 | 36.30±6.16 | 1.26 | 19.72±2.89 | 0.76 | 35.30 | 0.80 |

| Cell line | Drug | | | | | | | | | |
|---|---|---|---|---|---|---|---|---|---|---|
| | Cabozantinib | | Trametinib | | Alpelisib | | LY294002 | | AT13148 | |
| | **$IC_{50}$** (μM) | **RF** | **$IC_{50}$** (μM) | **RF** | **$IC_{50}$** (μM) | **RF** | **$IC_{50}$** (μM) | **RF** | **$IC_{50}$**(μM) | **RF** |
| HCC4006 | 8.52±1.44 | n/a | 0.017±0.005 | n/a | 2.84±0.46 | n/a | 12.61±3.00 | n/a | 2.30 | n/a |
| HCC4006$^r$ERLO$^{1\mu M}$ | 6.58±0.80 | 0.77 | 0.105±0.004 | 6.30 | 7.51±1.93 | 2.64 | 8.41±0.62 | 0.67 | 6.67 | 2.90 |
| HCC4006$^r$GEFI$^{1\mu M}$ | 8.73±1.62 | 1.02 | 0.043±0.007 | 2.58 | 8.27±1.44 | 2.91 | 13.03±2.88 | 1.03 | 15.24 | 6.63 |
| HCC4006$^r$AFA$^{100nM}$ | 2.48±0.64 | 0.29 | 0.073±0.010 | 4.36 | 4.15±1.14 | 1.46 | 3.65±0.53 | 0.29 | 11.34 | 4.94 |

**Supplementary Table 5. Determination of the $IC_{25}$ and $IC_{50}$ values fordifferent kinase inhibitors in HCC827, HCC4006, and their EGFR tyrosine kinase inhibitor-resistant sublines to different kinase inhibitors.** $IC_{50}$ values represent means of at least three biological repeats ± S.D. Drug response was determined by MTT assay after a 120h incubation period. $IC_{50}$ values were calculated using Calcusyn (Version 1.1, Biosof 1996).

| Cell line | Drug | | | | | | | |
|---|---|---|---|---|---|---|---|---|
| | Cabozantinib (μM) | | Trametinib (μM) | | Alpelisib (μM) | | LY294002 (μM) | |
| | **$IC_{25}$** | **$IC_{50}$** | **$IC_{25}$** | **$IC_{50}$** | **$IC_{25}$** | **$IC_{50}$** | **$IC_{25}$** | **$IC_{50}$** |
| HCC827 | **0.037** | **0.062** | **0.009** | **0.035** | **4.188** | **9.454** | **7.419** | **13.539** |
| HCC827$^{r}$ERLO$^{2μM}$ | **0.003** | **0.009** | **0.010** | **0.061** | **3.895** | **6.815** | **3.787** | **7.791** |
| HCC827$^{r}$GEFI$^{2μM}$ | **3.043** | **6.026** | **11.304** | **29.486** | **23.508** | **36.303** | **11.013** | **19.720** |
| HCC4006 | **3.226** | **6.580** | **0.015** | **0.105** | **1.463** | **7.511** | **1.235** | **8.405** |
| HCC4006$^{r}$ERLO$^{1μM}$ | **2.963** | **8.728** | **0.012** | **0.043** | **1.413** | **8.269** | **4.453** | **13.029** |
| HCC4006$^{r}$GEFI$^{1μM}$ | **0.018** | **2.479** | **0.022** | **0.073** | **0.755** | **4.148** | **1.000** | **3.648** |

**Supplementary Table 6: Impact of different kinase inhibitors on the sensitivity of EGFR tyrosine kinase-adapted HCC827 and HCC4006 sublines to their respective drugs of adaptation.** $IC_{50}$ values represent means of at least three biological repeats ± S.D. Drug response was determined by MTT assay after a 120h incubation period. $IC_{50}$ values were calculated using Calcusyn (Version 1.1, Biosof 1996). Resistance factors (RF) were calculated as follows $IC_{50}$ drug-resistant subline/ $IC_{50}$ respective parental cell line.

| Cell line | Untreated | | Drug ($IC_{25}$ treated) | | | | | | | |
|---|---|---|---|---|---|---|---|---|---|---|
| | | | Cabozantinib | | Trametinib | | Alpelisib | | LY294002 | |
| | **$IC_{50}$** (µM) | **RF** | **$IC_{50}$** (µM) | **RF** | **$IC_{50}$** (µM) | **RF** | **$IC_{50}$** (µM) | **RF** | **$IC_{50}$** (µM) | **RF** |
| HCC827 | 20.77±6.21 | 266.6 | 0.17 ± 0.06 | 2.2 | 3.11 ± 1.32 | 39.9 | 6.24± 0.73 | 80.1 | 5.97±1.52 | 76.6 |
| HCC827$^{r}$ERLO$^{2\mu M}$ | 8.16± 1.33 | 108.8 | 5.24 ± 2.48 | 69.8 | 6.09 ± 2.53 | 81.2 | 2.63± 0.91 | 35.0 | 5.10±2.17 | 68.0 |
| HCC827$^{r}$GEFI$^{2\mu M}$ | 1.74± 0.16 | 1319.5 | 0.44 ± 0.25 | 335.8 | 0.37± 0.15 | 282.1 | 0.0014±0.0008 | 1.0 | 0.003±0.0019 | 2.5 |
| HCC4006 | 18.17±1.84 | 691.9 | 35.59± 8.81 | 1355.4 | 29.88±8.77 | 1138.0 | 27.05 ± 18.39 | 1030.3 | 21.94 ± 13.05 | 835.5 |
| HCC4006$^{r}$ERLO$^{1\mu M}$ | 45.97±4.14 | 1569.5 | 6.68 ± 1.91 | 228.1 | 47.77±9.34 | 1631.0 | 16.06 ± 2.45 | 548.4 | 51.58 ± 11.13 | 1761.1 |
| HCC4006$^{r}$GEFI$^{1\mu M}$ | 2.24± 1.27 | 14807.9 | 3.28 ± 0.55 | 21645.7 | 2.81±1.67 | 18529.5 | 3.10 ± 0.73 | 20468.4 | 4.08 ± 1.74 | 26886.9 |
| Cell line | **Untreated** | | **Drug ($IC_{50}$ treated)** | | | | | | | |
| | | | Cabozantinib | | Trametinib | | Alpelisib | | LY294002 | |
| | **$IC_{50}$** (µM) | **RF** | **$IC_{50}$** (µM) | **RF** | **$IC_{50}$** (µM) | **RF** | **$IC_{50}$** (µM) | **RF** | **$IC_{50}$** (µM) | **RF** |
| HCC827 | 20.77±6.21 | 266.5 | 0.016± .008 | 0.2 | 2.29± 1.54 | 29.3 | 23.29 ± 4.44 | 298.9 | 22.97 ± 4.80 | 294.8 |
| HCC827$^{r}$ERLO$^{2\mu M}$ | 8.16 ± 1.33 | 108.8 | 5.80 ± 1.79 | 77.3 | 3.44± 0.56 | 45.8 | 1.83 ± 0.34 | 24.4 | 4.51 ± 2.20 | 60.1 |
| HCC827$^{r}$GEFI$^{2\mu M}$ | 1.74 ± 0.16 | 1319.5 | 0.03 ± 0.02 | 20.1 | 0.14± 0.12 | 104.9 | 0.0008±0.0007 | 0.6 | 0.026 ± 0.007 | 19.4 |
| HCC4006 | 18.17±1.84 | 691.9 | 30.07± 9.56 | 1145.3 | 52.06±23.72 | 1982.5 | 23.70 ± 0.62 | 902.7 | 36.78 ± 3.74 | 1400.8 |
| HCC4006$^{r}$ERLO$^{1\mu M}$ | 45.97±4.14 | 1569.5 | 2.31 ± 1.42 | 78.9 | 46.34± 3.04 | 1582.4 | 21.61 ± 5.36 | 737.8 | 23.97 ± 1.13 | 818.5 |
| HCC4006$^{r}$GEFI$^{1\mu M}$ | 2.24 ± 1.27 | 14807.9 | 1.22 ± 0.78 | 8065.6 | 2.19± 2.16 | 14418.0 | 3.57 ± 0.93 | 23555.4 | 3.40 ± 0.67 | 22455.5 |

**Supplementary Table 7. HDAC inhibitor concentrations inhibit HDACs with some level of specificity.**

| HDAC inhibitor | Concentration | HDACs inhibited | Reference |
|---|---|---|---|
| Apicidin | 200nM | HDAC (1)/2/3 | Bantscheff et al., 2011 |
| Bufexamac | 30µM | HDAC 6/10 | Bantscheff et al., 2011 |
| CI-994 | 2µM | HDAC 1/2/3/(8) | Bantscheff et al., 2011 |
| Compound 2 | 40µM | HDAC 8 | Krennhrubec et al., 2007<br>Oehme et al., 2009 |
| Fimepinostat/ CUDC-994 | 5nM | HDAC 1/2/3/10 | https://www.selleckchem.com/products/pi3k-hdac-inhibitor-i.html |
| Droxinostat | 15µM | HDAC 6/8 | https://www.selleckchem.com/products/Droxinostat.html |
| Mocetinostat | 300nM | HDAC 1/(2)/(3)/(11) | https://www.selleckchem.com/products/MGCD0103(Mocetinostat).html |
| Entinostat/ MS-275 | 250nM | HDAC 1(2)/3 | Deubzer et al., 2008<br>Witt et al., 2009 |
| Panobinostat | 10nM | pan | Witt et al., 2009 |
| PCI-24781 | 500nM | pan | Bantscheff et al., 2011 |
| PCI-34051 | 4µM | HDAC 8 | Balasubramanian et al., 2008<br>Bantscheff et al., 2011 |
| Romidepsin/ FK-228 | 40nM | HDAC 1/2 | Bantscheff et al., 2011 |
| Trichostatin A | 75nM | pan | Deubzer et al., 2008<br>Bantscheff et al., 2011 |
| Tubacin | 2.5µM | HDAC 6 | Witt et al., 2009 |
| Tubastatin A | 7.5µM | HDAC 6/10 | Géraldy et al., 2019 |
| Vorinostat/ SAHA | 1µM | pan | Bantscheff et al., 2011 |

**Supplementary Table 8. Effect of HDAC inhibitors on the viability of the project cell lines.** Values represent means of at least three biological repeats ± S.D. Drug response was determined by MTT assay after a 120h incubation period. $IC_{50}$ values were calculated using Calcusyn (Version 1.1, Biosof 1996).

| HDAC inhibitor | Cell viability (%) | | | | | | | |
|---|---|---|---|---|---|---|---|---|
| | HCC827 | | | | HCC4006 | | | |
| | HCC827 | $^{r}$ERLO$^{2μM}$ | $^{r}$GEFI$^{2μM}$ | $^{r}$AFA$^{50nM}$ | HCC4006 | $^{r}$ERLO$^{1μM}$ | $^{r}$GEFI$^{1μM}$ | $^{r}$AFA$^{100nM}$ |
| **Abexinostat** | 76% ± 17 | 44% ± 17 | 68% ± 17 | 71% ± 7 | 44% ± 5 | 48% ± 10 | 65% ± 28 | 40% ± 15 |
| **Panobinostat** | 78% ± 7 | 79% ± 8 | 97% ± 3 | 55% ± 12 | 56% ± 5 | 62% ± 8 | 77% ± 24 | 64% ± 24 |
| **TSA** | 82% ± 10 | 90% ± 7 | 91% ± 6 | 63% ± 16 | 72% ± 11 | 71% ± 7 | 75% ± 14 | 71% ± 21 |
| **Vorinostat/ SAHA** | 89% ± 10 | 56% ± 6 | 75% ± 12 | 66% ± 4 | 43% ± 2 | 47% ± 18 | 60% ± 23 | 47% ± 20 |
| **Apicidin** | 61% ± 13 | 28% ± 7 | 9% ± 11 | 70% ± 30 | 25% ± 13 | 41% ± 17 | 54% ± 24 | 36% ± 13 |
| **CI-994** | 82% ± 10 | 73% ± 5 | 88% ± 10 | 55% ± 11 | 39% ± 28 | 49% ± 43 | 52% ± 49 | 49% ± 29 |
| **CUDC -994** | 97% ± 4 | 85% ± 10 | 74% ± 27 | 64% ± 11 | 53% ± 14 | 58% ± 17 | 72% ± 34 | 61% ± 18 |
| **Entinostat/ MS-275** | 99% ± 1 | 71% ± 8 | 87% ± 12 | 66% ± 6 | 55% ± 26 | 68% ± 32 | 71% ± 29 | 68% ± 23 |
| **Mocetinostat** | 98% ± 4 | 49% ± 6 | 61% ± 16 | 62% ± 4 | 51% ± 9 | 59% ± 6 | 72% ± 24 | 59% ± 18 |
| **Tubacin** | 100% ± 0 | 91% ± 4 | 67% ± 14 | 74% ± 9 | 60% ± 12 | 71% ± 15 | 68% ± 6 | 64% ± 18 |
| **Bufexamac** | 92% ± 5 | 71% ± 11 | 80% ± 17 | 77% ± 11 | 32% ± 25 | 36% ± 33 | 48% ± 46 | 42% ± 25 |
| **Tubastatin A** | 92% ± 9 | 81% ± 3 | 89% ± 7 | 63% ± 17 | 66% ± 10 | 76% ± 17 | 85% ± 18 | 72% ± 21 |
| **Droxinostat** | 85% ± 5 | 62% ± 7 | 57% ± 18 | 62% ± 9 | 26% ± 19 | 30% ± 27 | 32% ± 29 | 25% ± 16 |
| **Cpd 2** | 87% ± 11 | 72% ± 4 | 65% ± 17 | 63% ± 14 | 33% ± 22 | 50% ± 16 | 56% ± 36 | 50% ± 19 |
| **PCI-34051** | 81% ± 8 | 78% ± 12 | 70% ± 18 | 74% ± 11 | 62% ± 6 | 63% ± 7 | 73% ± 13 | 64% ± 20 |
